# Inference of alveolar capillary network connectivity from blood flow dynamics

**DOI:** 10.1101/2024.01.22.576641

**Authors:** Kerstin Schmid, Andy L. Olivares, Oscar Camara, Wolfgang M. Kuebler, Matthias Ochs, Andreas C. Hocke, Sabine C. Fischer

## Abstract

The intricate structure of the lungs is essential for the gas exchange within the alveolar region. Despite extensive research on the pulmonary vasculature, there are still unresolved questions regarding the connection between capillaries and the vascular tree. A major challenge is obtaining comprehensive experimental data that integrates morphological and physiological aspects.

We propose a computational approach that combines data-driven 3D morphological modeling with computational fluid dynamics simulations. This method enables investigating the connectivity of the alveolar capillary network with the vascular tree based on the dynamics of blood flow. We developed 3D sheet-flow models to accurately represent the morphology of the alveolar capillary network and conducted computational fluid dynamics simulations to predict flow velocities and pressure distributions.

Our approach focuses on leveraging functional features to identify the most plausible architecture of the system. For given capillary flow velocities and arteriole-to-venule pressure drops, we deduce details about arteriole connectivity. Preliminary connectivity analyses for non-human species indicate that their alveolar capillary network of a single alveolus is linked to at least two arterioles with diameters of 20 µm or a single arteriole with a minimum diameter of 30 µm.

Our study provides insights into the structure of the pulmonary microvasculature by evaluating blood flow dynamics. This inverse approach represents a new strategy to exploit the intricate relationship between morphology and physiology, applicable to other tissues and organs. In the future, the availability of experimental data will play a pivotal role in validating and refining the hypotheses analyzed with our computational models.

**New and noteworthy:** The alveolus is pivotal for gas exchange. Due to its complex morphology and dynamic nature, structural experimental studies are challenging. Computational modeling offers an alternative. We developed a databased 3D model of the alveolar capillary network and performed blood flow simulations within it. Choosing a novel perspective, we inferred structure from function. We systematically varied properties of vessels connected to our capillary network and compared simulation results with experimental data to obtain plausible vessel configurations.

## 1 Introduction

Morphology and physiology are closely intertwined for organ function. In the lung, the minimal functional unit, the alveolus, is structured and composed for the vital exchange of carbon dioxide and oxygen between the air space and capillary blood system. This demand requires maximizing surface area while minimizing barrier thickness. Additionally, efficient blood flow is essential, necessitating minimal resistance to flow but allowing sufficient contact time for effective gas exchange at the same time.

Despite the inherent relationship between structure and function, experimental studies often consider them separately. Traditional morphological studies (Gehr et al., 1978; Weibel et al., 1993; Mühlfeld et al., 2010) rely on invasive techniques for organ fixation, often causing tissue distortion and compromising accuracy (Tuğcu et al., 2013; Braber et al., 2010; Hausmann et al., 2004; Javed et al., 1994; Weibel, 1984; Mooi and Wagenvoort, 1983). Physiology research on the whole lung scale can employ non-invasive methods (e.g. Bosman et al. (1991); Folke et al. (2003)). Yet, investigating processes of the alveolar compartment, such as blood flow distribution, remains a major challenge.

Computational modeling offers a unique avenue to integrate morphological and physiological research. Typically, *in silico* models aim to build accurate anatomical representations to predict organ function. In pulmonary research, this has included the development of increasingly sophisticated 2D and 3D models of the pulmonary microcirculation to predict the distribution of blood flow (Dhadwal et al., 1997; Huang et al., 2001; Burrowes et al., 2004). These approaches have primarily focused on the impact of morphological details on physiological output.

In this work, we demonstrate how physiological parameters can provide morphological insight. Our approach combines 3D morphological modeling of an alveolus and the alveolar capillary network (ACN) with blood flow simulations using computational fluid dynamics (CFD). Our focus is on uncharted morphological aspects of the ACN’s connection to the vascular tree. We link the 3D ACN model to generic arteriole and venule models, varying their numbers and diameters. We demonstrate how the choice of boundary conditions and model parameters such as the fluid viscosity influence the CFD simulation results, including capillary flow velocity and arteriole-to-venule pressure drop. With simulation settings based on findings from disparate experimental studies, our model produces results that are within the range of literature reference values. Additionally, we illustrate how the physiological parameters - capillary flow velocity and pressure drop - can be used to infer structural details about the linkage between the ACN and arterioles or venules. It is crucial to emphasize that the success of our approach relies on the availability of appropriate experimental data for validation.

## 2 Methods

For this integrative study, a number of specialized software tools were used in a multi-step workflow (Figure 1). We created 3D models of the ACN based on morphometric data from the literature. These models served as templates for building high-quality finite-element meshes, which were subsequently used to run CFD simulations to predict blood flow dynamics.

**Figure 1:**
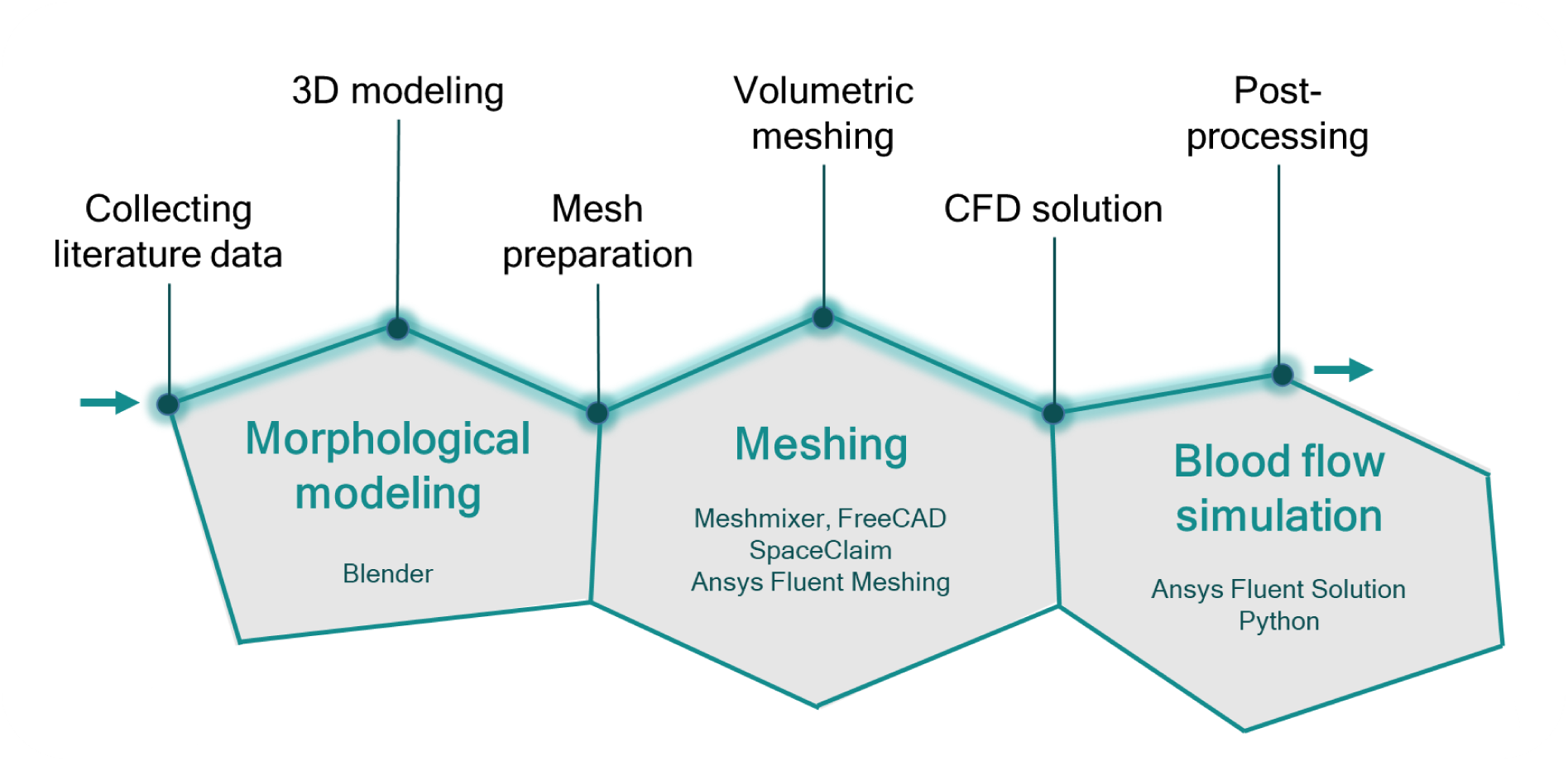
The methodological concept of this work combines morphological *in silico* modeling of the alveolus and the alveolar capillary network with blood flow simulations. Relevant morphological data was collected from existing literature. Using the software Blender, 3D models of the alveolus and the alveolar capillary network were generated. These models underwent mesh preparation using Meshmixer, FreeCAD, and SpaceClaim to ensure compatibility with the Ansys software. Ansys Fluent Meshing was employed to create volumetric meshes for computational fluid dynamics simulations. The simulations were conducted with Ansys Fluent Solution. Post-processing, including data extraction and analysis, was performed using Ansys Fluent Solution and Python scripting.

### 2.1 Morphological modeling

The true-to-scale, three-dimensional models of the capillary network around a single alveolus were manually created with the modeling software Blender v.3.3 (Community, 2018). Emphasis was placed on ensuring that the scale and size ratios correspond to quantitative morphological data found in the literature.

#### 2.1.1 Alveolar base

Initially, a basic geometry of an alveolus was created, which will be referred to as “alveolar base” in the following. It was constructed as an open 3/4 spheroid with an opening, the so called alveolar mouth (Figure 2 A). The diameter or volume of the spheroid, as well as its depth and the diameter of the opening were based on published stereological measurements, making them the input parameters for our modeling process. On the other hand, we employed a separate set of parameters for evaluation, comparing them with reference values from the literature. These “output parameters” encompass the surface area and volume or diameter of the alveolar base. All parameter values are listed in Table 1 of the Results section.

**Figure 2:**
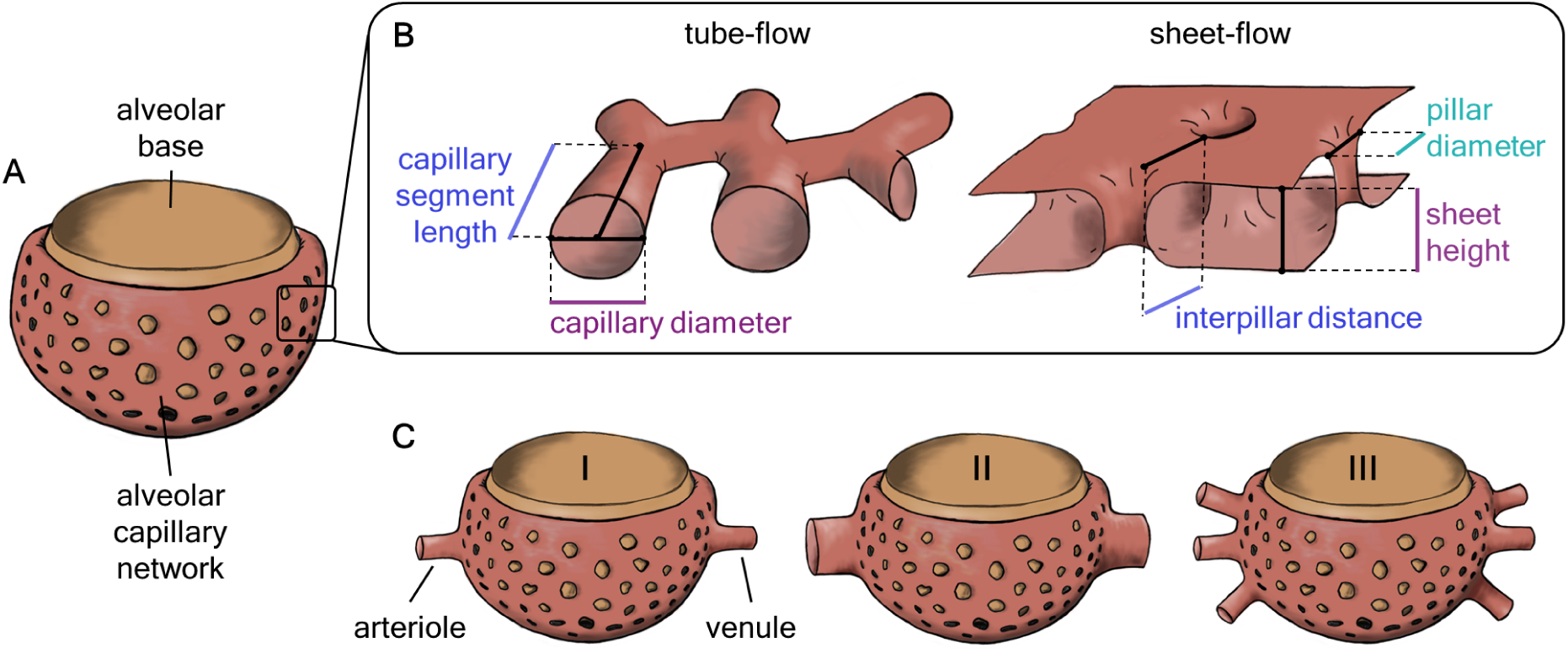
A) The alveolar capillary network is built around an open 3/4 spheroid, the alveolar base. B) Two different concepts of the capillary bed geometry, the tube-flow and the sheet-flow, were compared. A tube-flow capillary network is parameterized by the length and diameter of the capillary segments. In contrast, the sheet-flow capillary bed is defined by the height of the sheet, the diameter of the tissue pillars and their distance from each other. C) The capillary network was connected to different configurations of arterioles and venules. Starting from the default configuration (I) of one arteriole and one venule, both 20 µm in diameter, either the diameter (II) or the number (III) of vessels was varied.

**Table 1:** Comparison of in- and output parameter values of 3D geometrical models *Alveolus_D_* and *Alveolus_V_* with morphometric measurements from the literature. The difference between the two models is that the input parameters for *Alveolus_D_* included the diameter and for *Alveolus_V_* the volume of the alveolus. By comparing the resulting output parameters with measurements from the literature, the models were evaluated. References: [1](Hansen and Ampaya, 1975), [2](Mercer et al., 1994), [3](Ochs et al., 2004), [4](Kawakami and Takizawa, 1987), [5](Toshima et al., 2004), [6](Matsuda et al., 1987), [7](Stone et al., 1992).

| Parameter | Literature | *Alveolus<sub>D</sub>*<br>Model | *Alveolus<sub>V</sub>*<br>Model |
| --- | --- | --- | --- |
| **Modeling (input)** |  |  |  |
| Alveolus - Shape | 3/4 spheroid, truncated cone, 1/4 spheroid, ...[1] | 3/4 spheroid | 3/4 spheroid |
| Alveolus - Diameter ($\mu\text{m}$) | 225 [2], 232 [1] | 225 | - |
| Alveolus - Volume ($10^6 \mu\text{m}^3$) | 4.6 [1], 4.2 [3], 3.4 [2] | - | 4.6 |
| Pores of Kohn - Number | 17 [4] | 17 | 17 |
| Pores of Kohn - Diameter ($\mu\text{m}$) | 7 - 19 [4], 7.2 - 16.5 [5] | 12.3 | 12.3 |
| Alveolar Mouth - Diameter ($\mu\text{m}$) | 181 [1], 220 [6] | 181 | 181 |
| **Evaluation (output)** |  |  |  |
| Alveolus - Diameter ($\mu\text{m}$) | 225 [2], 232 [1] | - | 216 |
| Alveolus - Volume ($10^6 \mu\text{m}^3$) | 4.6 [1], 4.2 [3], 3.4 [2] | 5.3 | - |
| Alveolus - Depth ($\mu\text{m}$) | 174 [1] | 178 | 167 |
| Alveolus - Surface Area ($10^5 \mu\text{m}^2$) | 1.2 [1,2], 2.1 [7] | 1.52 | 1.39 |
| Alveolar Mouth - Area ($10^5 \mu\text{m}^2$) | 0.3 [1] | 0.26 | 0.24 |

#### 2.1.2 Alveolar capillary network

The capillary network was built around this alveolar base. We considered two alternative concepts of the morphology of the alveolar capillary bed that have been proposed in the literature, the tube-flow (Mühlfeld et al., 2010) and the sheet-flow (Fung and Sobin, 1969), as shown in Figure 2 B. In the tube-flow model, the capillary bed is considered as a network of cylindrical tubes with a defined diameter and length. In the sheet- flow model, the capillary bed is regarded as a continuous sheet formed by two endothelial layers. These layers are held apart by connective tissue and cellular pillars at a specific height, i.e., the thickness of the sheet. The diameter and spacing of the pillars further determine the structure of the capillary bed.

We followed different modeling approaches for the tube-flow and the sheet-flow models of the ACN, based on correspondingly different input parameters. For the tube-flow model, the diameter and length of capillary segments were aligned with measurements from Mühlfeld et al. (2010). The sheet-flow model was built taking into account the height of the sheet, the diameter of the tissue pillars and their distance from each other. Existing literature only provides measurements of cats for these parameters (Sobin et al., 1970). In order to create a sheet-flow model that matches the morphological proportions of the human ACN, the following assumptions were made: The height of the capillary sheet corresponds to the diameter of capillary tubes and the distance between pillars corresponds to the capillary segment length. The pillar diameter was derived by assuming the same ratio between pillar diameter and interpillar distance as observed in cats, roughly 0.5 (Sobin et al., 1970). All parameter values are listed in Table 2 of the Results section. Further details on the modeling workflow in Blender are provided in Supplementary Section S.1. For evaluation purposes, the volume and surface area of the 3D ACN models were compared with reference values from morphological studies.

**Table 2:** *In silico* models of the human alveolar capillary network (ACN) have lower surface area and volume compared to corresponding estimates from morphometric studies. Tube-flow and sheet-flow models of the ACN around the *Alveolus_D_* and *Alveolus_V_*models, respectively, were built based on literature reference values for the input parameters. Output parameters were measured after the modeling process and compared with literature values for evaluation purposes. References: [7](Stone et al., 1992), [8](Mühlfeld et al., 2010), [9](Gehr et al., 1978). *Derived from interpillar distance assuming a ratio between pillar diameter and interpillar distance of 0.5. **Measurements for the entire lung were converted to an alveolus, assuming 480 million alveoli in the human lung (Ochs et al., 2004).

| Parameter | Literature | *Alveolus<sub>D</sub>* Model |  | *Alveolus<sub>V</sub>* Model |  |
| --- | --- | --- | --- | --- | --- |
|  |  | sheet-flow | tube-flow | sheet-flow | tube-flow |
| Modeling (input) |  |  |  |  |  |
| Capillary diameter / sheet height (μm) | 6.3 [8] | 6.34 | 6.31 | 6.36 | 6.3 |
| Capillary segment length / interpillar distance (μm) | 5.9 [8] | 6.17 | 5.75 | 6.33 | 5.54 |
| Capillary pillar diameter (μm) | 3* | 3.1 | - | 3.23 | - |
| Evaluation (output) |  |  |  |  |  |
| Capillary volume (10<sup>5</sup> μm<sup>3</sup>) | 8.2** [8],<br>8.8** [9] | 7.12 | 6.09 | 6.32 | 5.54 |
| Capillary surface area (10<sup>5</sup> μm<sup>2</sup>) | 5.4** [8],<br>5.3** [9],<br>3.0** [7] | 3.42 | 3.23 | 3.12 | 2.93 |

#### 2.1.3 Connectivity models

We set out to investigate the linkage of the ACN to the rest of the pulmonary vascular tree. There is no distinct perfusion unit as the ACN is a continuum connected to multiple arterioles and venules and spanning multiple alveoli (Buchacker et al., 2019). To characterise the tissue structure, it is therefore useful to determine the ratio of the number of arterioles and venules to alveoli. Specifications for the diameter of arterioles and venules in the literature range from 13 to 30 µm (Horsfield, 1978; Horsfield and Gordon, 1981; Pump, 1966; Huang et al., 1996). In addition, it was also observed how alveolar capillaries branch off from rather large arteries with diameters of up to 100 µm (Horsfield, 1978). Based on this information, we created different configurations of arterioles and venules connected to our ACN model for the connectivity analyses (Figure 2 C). The default configuration was one arteriole and one venule, both with a diameter of 20 µm and positioned opposite each other. Starting from this, we created two different sets of models. In the first set, we varied the diameter of the vessels connected to the ACN from 20 µm to 60 µm in increments of 10 µm. This was done either symmetrically or only for one of the vessel types (arteriole or venule) at a time. In the second set, the number of connected arterioles and venules was increased from one to five. Again, the changes were made both symmetrically and for each vessel type individually.

### 2.2 Volumetric mesh preparation

The models of the ACN created in Blender were hollow bodies with their surface described by an average of 2.1*·*10^6^ nodes and faces and 4.2*·*10^6^ edges. For blood flow simulations using CFD, the fluid domain, i.e., the volume inside the capillary, must be defined by a discrete volumetric mesh. Hence, a finite volume (FV) mesh combines both components: A surface mesh, describing the wall of the geometry as well as inlet and outlet faces and the volumetric mesh defining the interior of the geometry (Figure 3). To be able to simulate the interaction of the fluid with the wall, a particularly high resolution of the mesh elements close to the wall is necessary. The volumetric mesh is therefore divided into a high-resolution boundary layer of prism elements near the wall and a coarser layer of polyhedral elements in the center.

**Figure 3:**
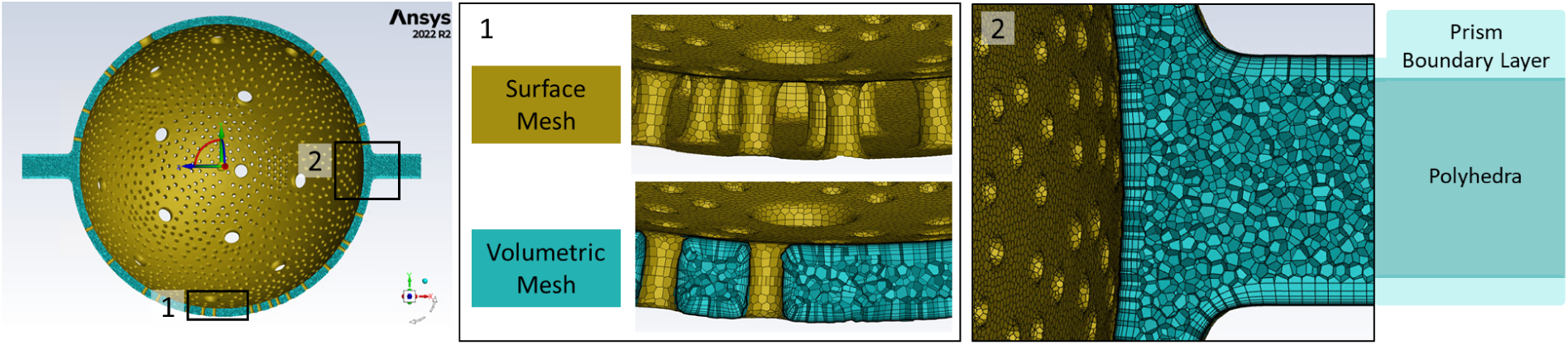
A finite volume mesh generated with the Ansys^®^ Fluent Meshing software. The surface mesh (gold) defines the capillary wall and inlet and outlet surfaces as they are given by the Blender model. The volumetric mesh (turquoise) consists of a boundary layer of prism elements near the wall and polyhedral elements inside the volume. The prism elements of the boundary layer increase in thickness from the wall towards the polyhedral elements.

In order to serve as a template for FV meshes, the Blender geometries of the ACN first had to be processed in several steps using different software programs. First, the geometries were exported from Blender as STL (stereolithography) files. In STL files, the surface of a 3D object is defined by a mesh of triangular faces. With Autodesk^®^ Meshmixer™version 3.5.474 (Inc., 2019), the resolution of this surface mesh was then reduced to roughly 500,000 triangles without distorting the geometry. In a next step, the STL files were converted to a CAD (computer aided design) file using the open-source software FreeCAD version 0.20.1 (FreeCAD, 2022). In this format, they were compatible with Ansys’ modeling software SpaceClaim, where the ACN geometry was organized into inlet, wall and outlet zones. Finally, the FV mesh was created with the Ansys Fluent Meshing software using the guided workflow “watertight geometry”. This is the primary workflow to generate FV meshes from clean boundary meshes, i.e., geometries without any leakage, as was the case with our geometries. More details on the meshing workflow are included in Supplementary Section S.2. The quality of the meshes was quantified in terms of the parameters inverse orthogonal quality, size change and aspect ratio (Supplementary Section S.3).

### 2.3 Computational fluid dynamics

We simulated blood flow in the morphometric models described in Section 2.1.3 using computational fluid dynamics (CFD). The CFD study was performed with the software Ansys^®^ Academic Research CFD, release 2022 R2 (ANSYS, 2022). A physical model describing the fluid flow was solved numerically in the domain of the ACN models (Section 2.3.1). We evaluated and estimated the quality of our FV meshes, as well as our CFD solution, and performed technical replications (Supplementary Section S.5).

#### 2.3.1 CFD solution

The three-dimensional flow of a fluid was described by a system of second order nonlinear partial differential equations based on the conservation of mass (1) and momentum (2) (ANSYS, 2009). The equation for the conservation of mass (or continuity equation) is expressed as:

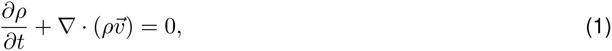

where ∇ is the divergence operator applied to both the density *ρ* and the velocity vector *v⃗*. In the case of an incompressible fluid, the rate of density change 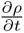 can be neglected. The conservation of momentum was expressed as:

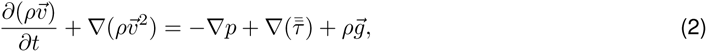

where *p* is the static pressure, 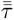 the stress tensor which depends on molecular viscosity and *ρg⃗* is the gravitational body force.

In Ansys Fluent Solution, these equations were represented as algebraic expressions in discrete domains of space (finite-volume method) and a pressure-based solver was used to solve them iteratively for a steady state. An estimation of the Reynolds number for the flow in our model resulted in 0.0076. Therefore, the viscous model was set to laminar flow. Vessel walls were assumed rigid, and a no slip boundary condition was applied at the walls. A detailed list of simulation settings made in Ansys is given in Supplementary Section S.4. The target values recorded were the mass-weighted average of flow velocity within capillaries and the pressure drop, calculated from the mass-weighted averages of static pressure at the inlet and outlet surfaces. Additionally, the global mass balance was monitored for every iteration during the simulation. Convergence criteria were met when the scaled and normalized residual values fell below the threshold of 1*·*10^-5^. Further, it was manually controlled that all target values had reached a constant value and that the global mass balance (defined as the sum of mass flow rates of all inlet and outlet faces) had leveled out around 0. More information on the convergence criteria are included in Supplementary Section S.5.

#### 2.3.2 Fluid properties and boundary conditions

Blood was approximated using two approaches: First, as a Newtonian fluid with a constant viscosity of 0.002 kg/(m*·*s). Secondly, as a non-Newtonian fluid using the Carreau viscosity model. This model describes the apparent viscosity *η* of a fluid as a function of shear rate *γ* as follows:

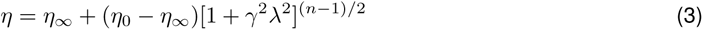

where *η*_0_ = 0.056*Pa · s* and *η_∞_* = 0.0035*Pa · s* are the viscosity values at zero and infinitely high shear rates, *λ* = 1.902s is a time constant and *n* = 0.3568 the power-law index. The parameter values appropriate for blood conditions were adopted from Albors et al. (2023).

Fluid density was set constant at 1050 kg/m^3^. Boundary conditions were set at 0.0023 m/s inlet velocity (2.3 mm/s (Horimoto et al., 1979)) and 1127 Pa outlet pressure (11.5 cmH_2_O (Nagasaka et al., 1984)).

#### 2.3.3 Post-processing

In Ansys, the results were visualized in the 3D model using pathlines or heatmaps. A pathline represents the path of a particle over time. They were color-coded according to the velocity magnitude. Heatmaps illustrated the distribution of static pressure in the model. Analysis of the quantitative simulation output and the presentation of the results by graphs was done in python v.3.9.12. Means and standard deviations of four technical replicates are given. Regression analyses were performed on the relationship between target values and arteriolar volume. To facilitate an easier comparison of our simulation results with literature values, we converted the units used in Ansys, which were m/s for velocity and Pa for pressure, to mm/s and mmHg, respectively.

#### 2.3.4 Sensitivity Analyses

Using the Newtonian fluid model, the boundary conditions inlet velocity and outlet pressure, as well as the fluid properties density and viscosity were varied separately while maintaining the standard settings used in our main experiments (Table S.3). Regarding the boundary conditions and fluid density, we selected value ranges of reasonable magnitude, with our standard values positioned approximately in the middle of these ranges. Specifically, we varied the inlet velocity from 0.5 mm/s to 3 mm/s in 0.25 mm/s increments, the outlet pressure from 500 Pa (3.75 mmHg) to 1500 Pa (11.25 mmHg) in 100 Pa increments and density between 500 kg/m^3^ and 1500 kg/m^3^ in 100 kg/m^3^ increments. The viscosity value range was chosen to encompass both the viscosity value of our Newtonian fluid model (2 cP) and all values applicable within the non-Newtonian viscosity model. It ranged from 1 cP to 56 cP in steps of 5.5 cP.

## 3 Results

### 3.1 Evaluation of 3D morphological models of alveolar capillary network

The first milestone of our work was a 3D model of the ACN around an alveolus based on published morphological measurements. When modeling the alveolar base, the following behavior was observed: If the model was based on the literature values for the diameters of the alveolus and the alveolar opening, the alveolar volume exceeded the respective measurement from the same study (Hansen and Ampaya, 1975) (Figure 4). Conversely, an alveolus model with the appropriate volume had a smaller diameter than specified in the literature (Table 1). For the time being, we continued with two different models, either based on the diameter (*Alveolus_D_*) or on the volume (*Alveolus_V_*).

**Figure 4:**
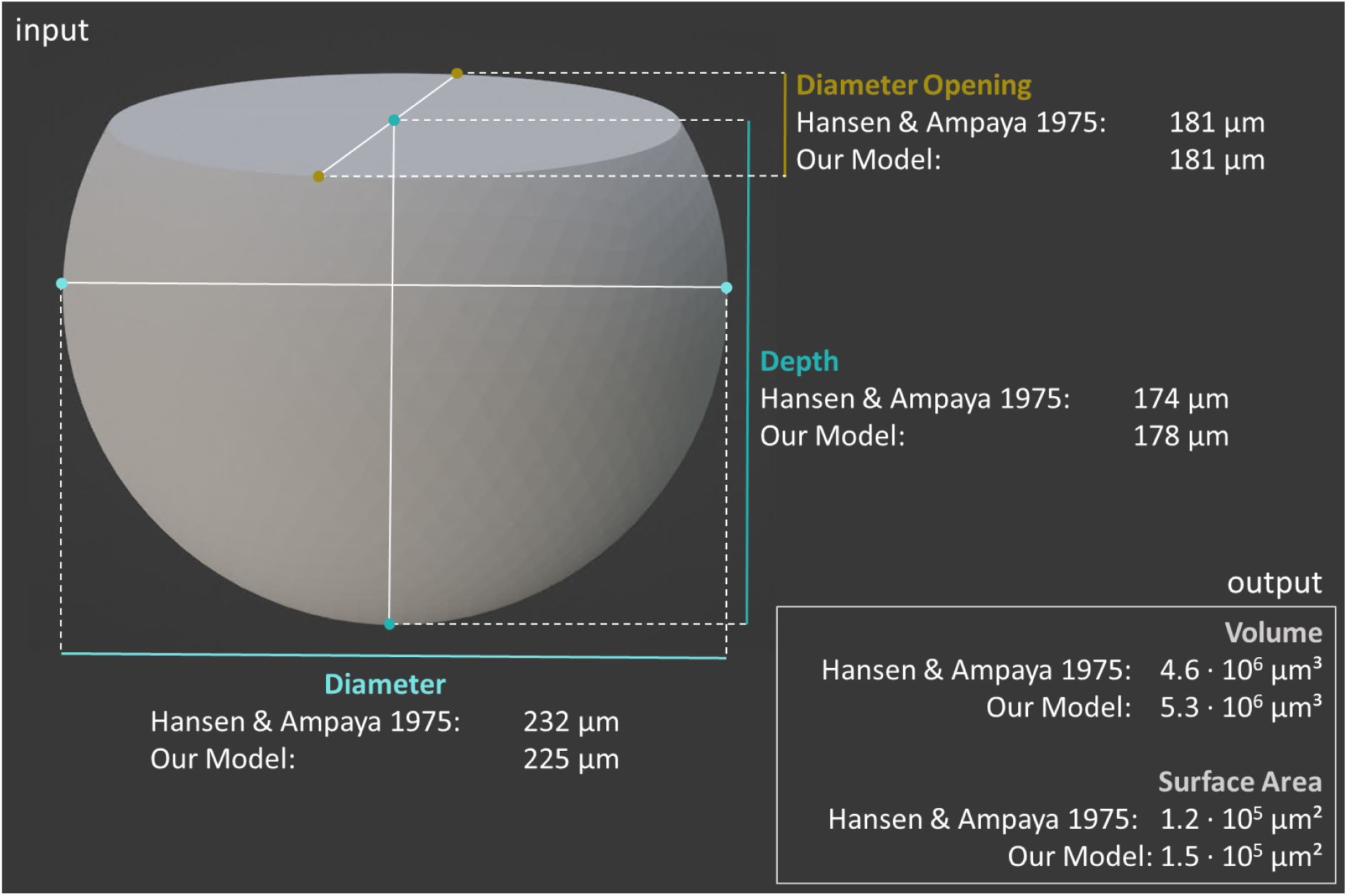
Comparison of in- and output parameter values of the 3D geometrical model of an alveolus with morphometric measurements from the literature (Hansen and Ampaya, 1975). The basis of the 3D model was an open spheroid for which the diameter, depth and diameter of the opening (input parameters) were matched to the respective literature values.

Next, we added the ACN to the *in silico* alveolus model. We compared tube-flow and sheet-flow models of capillary beds around the *Alveolus_D_* and *Alveolus_V_* base models (Table 2). The volume and surface area of these models were found to be lower than the values reported in the literature. In general, however, the *Alveolus_D_* ACN models were closer to the literature values than the *Alveolus_V_* ACN models and the same tendency applied to the sheet-flow models compared to the tube-flow models. Since the *Alveolus_D_* sheet-flow model also easier to implement, we considered it the best candidate for further work.

### 3.2 Blood flow simulations in a 3D model of the alveolar capillary network

Next, we used CFD simulations to predict blood flow in our 3D sheet-flow model of the ACN. The first step of this CFD study comprised sensitivity analyses to evaluate the effect of variations in boundary conditions at the inlet and outlet, and in the fluid properties of density and viscosity, on the simulation results. The flow velocity at any point in the model depends on the mass flow rate and the fluid density (Equation 1). The mass flow rate is determined by the inlet velocity boundary condition. Accordingly, a linear relationship could be observed between the inlet velocity and the mean velocity within capillaries (Figure 5). The static pressure at the outlet and the viscosity of the fluid had no effect on this simulation result. Similarly, the range of density values tested in this analysis had no effect on the flow velocity within capillaries, nor on the pressure drop from arteriole to venule (data not shown). For the calculation of the static pressure, the conservation of momentum (Equation 2) must be considered. This includes a dependence on viscosity. Correspondingly, the pressure drop showed a linear dependence on both inlet velocity and fluid viscosity (Figure 6). Although changes in the static pressure at the outlet affected the overall pressure in the model, the pressure drop remained unchanged.

**Figure 5:**
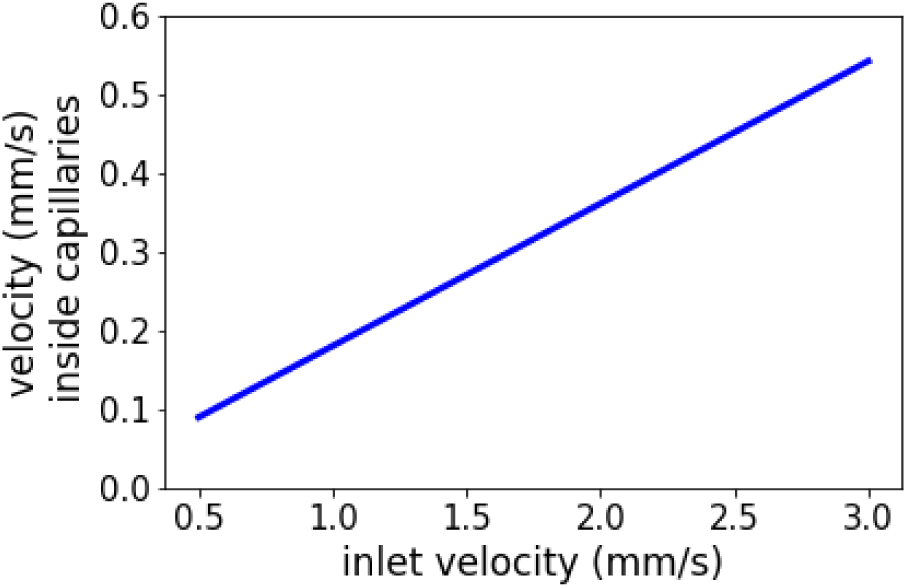
Sensitivity analysis for the effect of inlet velocity on the target value mean flow velocity within capillaries. Simulations were performed in the default model with one arteriole and one venule, both 20 µm in diameter. The inlet velocity was varied from 0.5 mm/s to 3 mm/s in steps of 0.25 mm/s.

**Figure 6:**
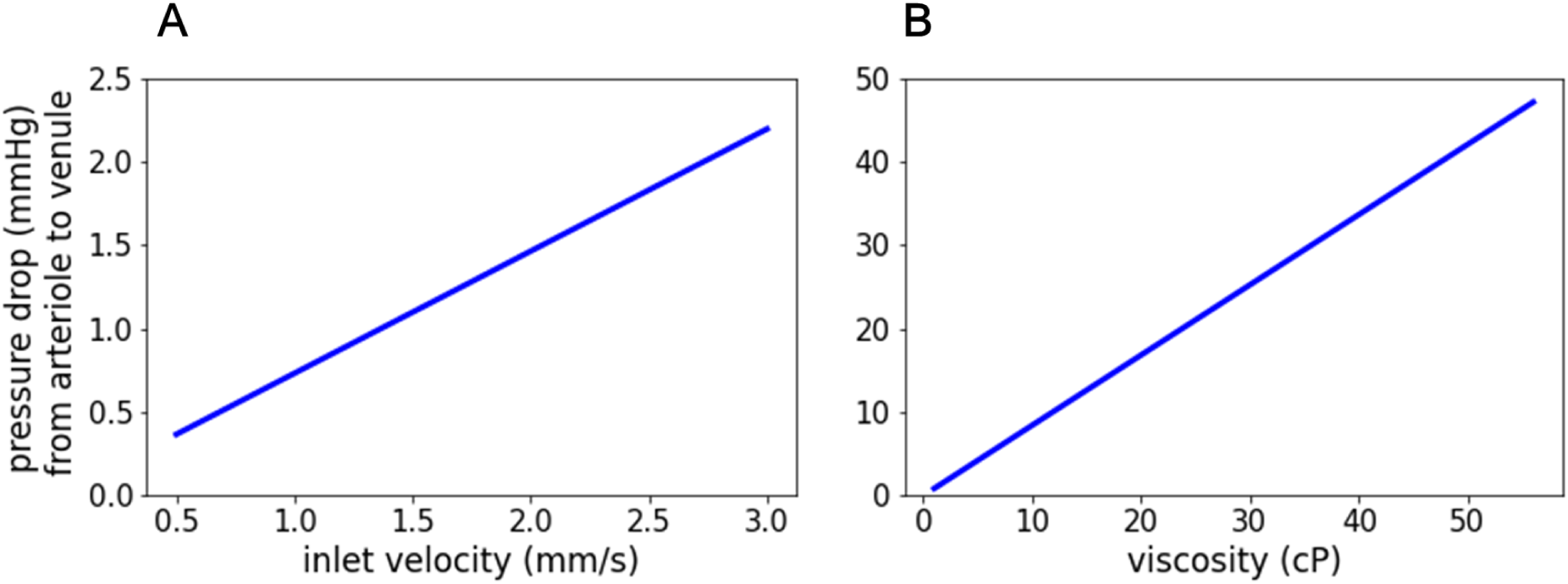
Sensitivity analyses for the effects of inlet velocity and fluid density on the target value pressure drop from arteriole to venule. Simulations were performed in the default model with one arteriole and one venule, both 20 µm in diameter. (A) The inlet velocity was varied from 0.5 mm/s to 3 mm/s in steps of 0.25 mm/s. (B) The viscosity was varied between 1 cP and 56 cP in steps of 5.5 cP.

With these dependencies in mind, we compared Newtonian and non-Newtonian viscosity models in the following experiments, keeping the fluid density and boundary conditions constant at their standard values (Supplementary Table S.3). Overall, the CFD simulations in our 3D ACN models showed that flow velocity inside the capillaries was lower compared to inside the inflow and outflow vessels (Figure 7). Additionally, the flow directions parallel to the orientation of these vessels were strongly preferred. This arrangement of arteriole, ACN and venule, resulted in some areas of the capillary net being more perfused than others. Regarding the mean flow velocity within the capillary network, the simulations with Newtonian and non-Newtonian viscosity models yielded consistent results of 0.39 mm/s *±*0.004 and 0.4 mm/s *±*0.003, respectively. In both cases, the pressure was evenly distributed across the model from the maximum at the inlet surface to the minimum at the outlet surface. However, the pressure drop in the Newtonian simulations, with a value of 1.64 mmHg *±*0.043, was markedly lower than the 3.54 mmHg *±*0.08 observed in the non-Newtonian simulations.

**Figure 7:**
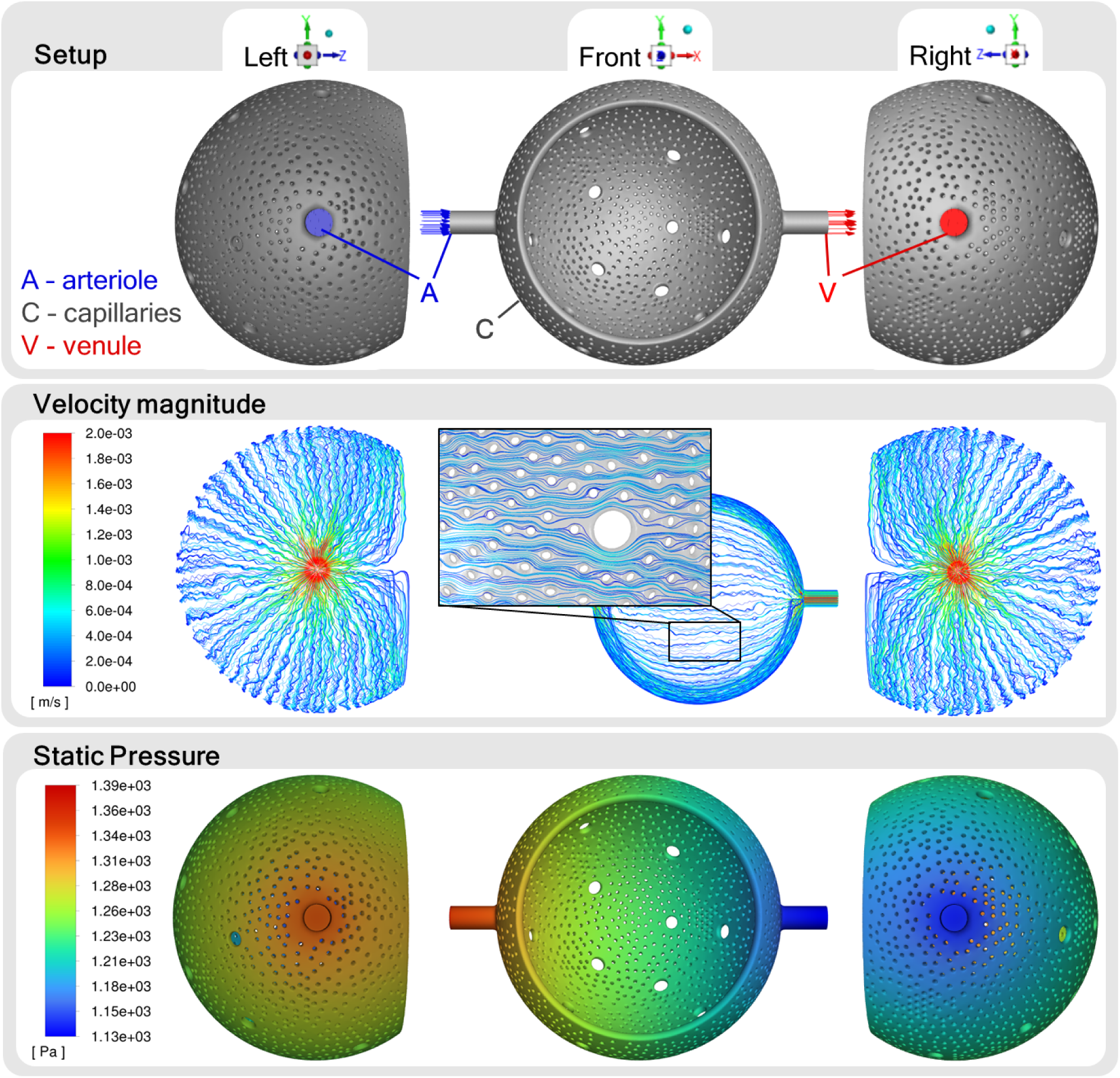
Results of computational fluid dynamics simulations in the 3D alveolar capillary network model. The capillary network was connected to one arteriole (inlet) and one venule (outlet), each with a diameter of 20 µm. Blood was approximated as a Newtonian fluid. Pathlines are color-coded according to the velocity magnitude, obtaining a mean flow velocity within the capillaries of 0.39 mm/s. The pressure decreased continuously from its maximum at 1345 Pa (10.09 mmHg) to its minimum at 1127 Pa (8.45 mmHg).

In summary, we extended our *in silico* model by including blood flow simulations. The flow velocity at the inlet has a positive linear effect on both target variables. The viscosity of the fluid has a positive linear effect on the pressure drop. Accordingly, differences in the pressure drop were observed in simulations with Newtonian and non-Newtonian viscosity models.

### 3.3 Inference of connection of the ACN to arterioles from physiological measurements

We investigated the linkage between the ACN and the pulmonary vascular tree by comparing models with different configurations of generic arterioles and venules. First, symmetrical models were considered in which arterioles and venules had the same configuration. To compare between configurations of vessel number and of diameter, we considered vessel volume as a common parameter. Both mean velocity within capillaries and pressure drop from arteriole to venule rose linearly with increasing volume of arterioles and venules (Figure 8). With respect to mean flow velocity, we did not observe differences in this relationship depending on whether the increase in volume was due to changes in vessel diameter or the number of vessels. The regression lines, expressed as v = *−*0.01 + 0.000032 *·* V and v = 0.03 + 0.000030 *·* V respectively, closely aligned. However, the pressure drop exhibited a more pronounced increase with vessel volume when the vessel diameter was enlarged (v = 0.83 + 0.00008 *·* V), as opposed to the increase resulting from a higher number of vessels (v = 1.01 + 0.00005 *·* V).

**Figure 8:**
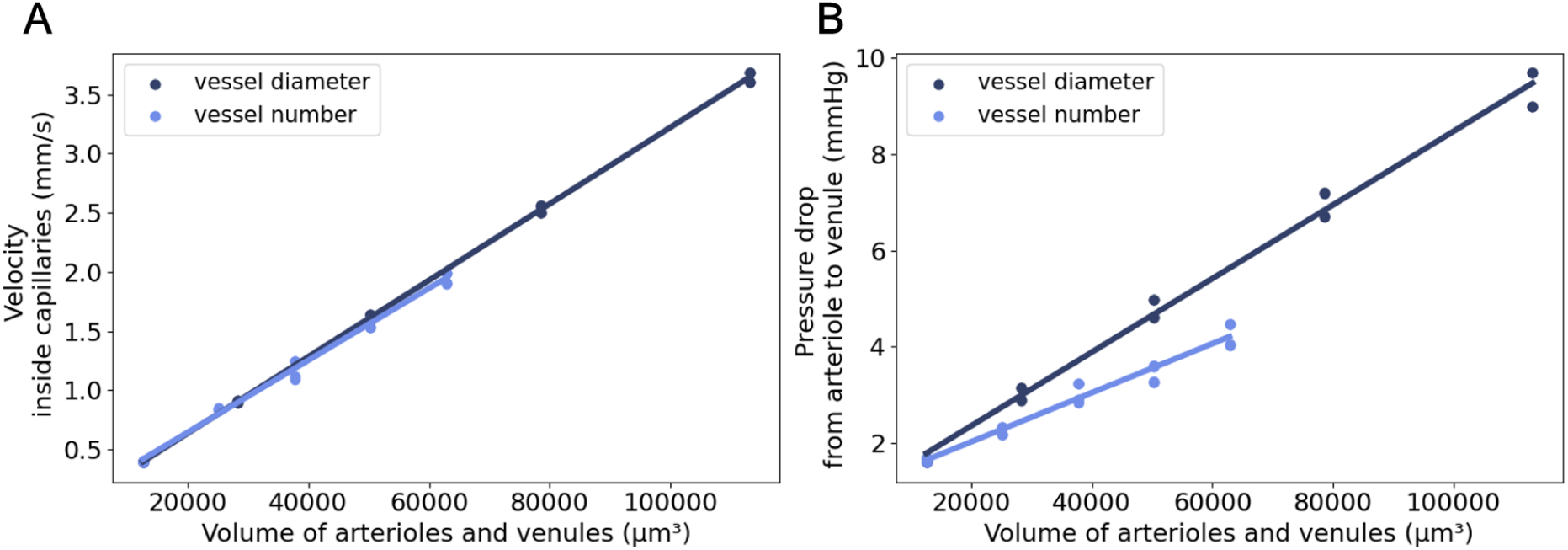
I*n silico* hemodynamic indices from flow simulations in symmetric connectivity models. The alveolar capillary network model was connected to either a growing number of arterioles and venules, ranging from one to five (light blue), or to one set of vessels with increasing diameters, ranging from 20 µm to 60 µm (dark blue). After computational fluid dynamics simulations of Newtonian fluid flow, the resulting mean velocity within capillaries (A) and pressure drop from arteriole to venule (B) were plotted against the volume of the inlet- and outlet vessels.

Next, we included asymmetric models where only one vessel type (arteriole or venule) was configured while the other remained in the default configuration. With increasing number or diameter of arterioles, both target values increased (Figure 9). The exact values are given in Supplementary Tables S.4 and S.5. Regarding the mean velocity within capillaries, we did not observe a difference between symmetrical and asymmetrical arteriolar configuration. The increase in pressure drop, however, was more pronounced in the asymmetric models compared to the symmetric models. Increasing the number or diameter of connected venules alone had no clear effect on the target values. Additional measurements at various positions in the model revealed that the asymmetric configuration of the venules only influenced the flow velocity within the venules, but not within the capillary network (Supplementary Figures S.3 and S.4).

**Figure 9:**
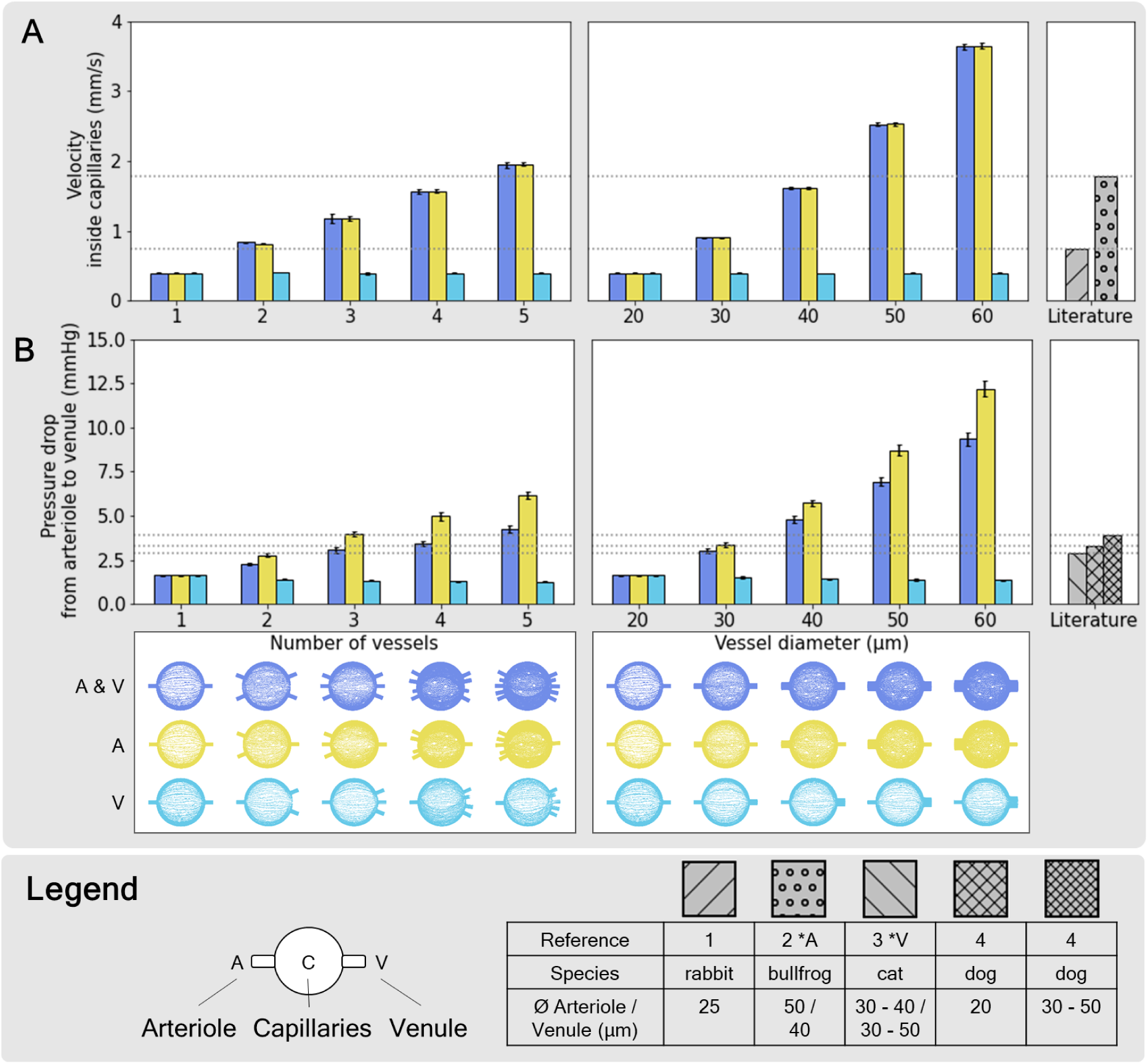
I*n silico* hemodynamic indices from flow simulations in symmetric and asymmetric connectivity models. Alveolar capillary network models were connected to increasing numbers (left panels) or diameters (right panels) of arterioles (yellow), venules (turquoise), or both (blue). Flow velocity within capillaries (A) and pressure drop from arteriole to venule (B) were predicted with computational fluid dynamics simulations using a Newtonian viscosity model. The results were compared with literature values of ^1^rabbit (Schlosser et al., 1965), ^2^bullfrog (Horimoto et al., 1979), ^3^cat (Nagasaka et al., 1984) and ^4^dog (Bhattacharya and Staub, 1980). **\***A Simulation boundary condition for inlet velocity at the arterioles were based on a reference from this publication. **\***V Simulation boundary condition for outlet pressure at the venules was based on references from this publication.

We compared our results with reference values from different species available in the literature (Schlosser et al., 1965; Horimoto et al., 1979; Nagasaka et al., 1984; Bhattacharya and Staub, 1980). These publications include information on the diameters of arterioles and venules (Figure 9 Legend), but the number of these vessels per alveolus is not specified. Our CFD simulation results using the Newtonian viscosity model aligned with the literature values for capillary velocity and arteriole-to-venule pressure drop, despite occasional discrepancies in arteriolar diameters. The comparison suggests that the ACN of a single alveolus is linked to at least arterioles with diameters of 20 µm or a single arteriole with a minimum diameter of 30 µm. In the results obtained from simulations using a non-Newtonian viscosity model, we made similar observations regarding velocity (Supplementary Figure S.6 A). However, the pressure drop was distinctly higher in these simulations (Supplementary Figure S.6 B), so that only models with a single 20-µm-arteriole provided results in the range of the literature values.

Further, we demonstrate how the regression of the simulation results can be used to infer the configuration of arterioles based on measurements of capillary velocity and pressure drop (Figure 10). Notably, the regression curves for capillary velocity overlap to the extent that distinctions between models with different vessel configurations become indistinguishable. In contrast, the regression curves for pressure drop yield identifiable variations. With these analyses at hand, we aimed to predict the arteriolar volume using a pair of example values. In the absence of available experimental data, we selected values from our simulation outcomes, opting for the scenario featuring each four arterioles and venules, all 20 µm in diameter. The corresponding simulation results were a capillary flow velocity of 1.56 mm/s and a pressure drop of 3.44 mmHg. According to the regressions, this velocity value is associated with a vessel volume of around 50,000 µm^3^ — equivalent to four vessels 20 µm in diameter or single 40-µm-diameter vessels (Figure 10 A). Regarding the pressure drop value, the distinct regression lines predict different possible vessel volumes. The regression analysis of the model set comprising symmetrical variations in the number of vessels points to a volume of 48,000 µm^3^. This closely corresponds to a model with four vessels, each 20 µm in diameter and thus aligns with the result obtained from the velocity regression. Additionally, this pressure drop value could have been measured in models with an arteriolar volume of less than 40,000 µm^3^, for example in the form of a single 30-µm-arteriole and a single 20-µm-venule. However, these models can be excluded as they do not agree with the velocity regression analysis. In conclusion, from the mean flow velocity within capillaries we can deduce the vessel volume. From the pressure drop regression, we obtain further information on possible configurations of arterioles and venules. In combination, we can identify the model geometry associated with this pair of values. We performed such an analysis with several pairs of values and in most cases obtained two alternative connectivity configurations (e.g. Supplementary Figure S.5), differing in whether the arteriolar volume is due to the vessel number or diameter. Hence, assuming that experimental measurements for capillary flow velocity and pressure drop from arteriole to venule would be available, we can make an unambiguous or two alternative predictions about the connectivity configuration.

**Figure 10:**
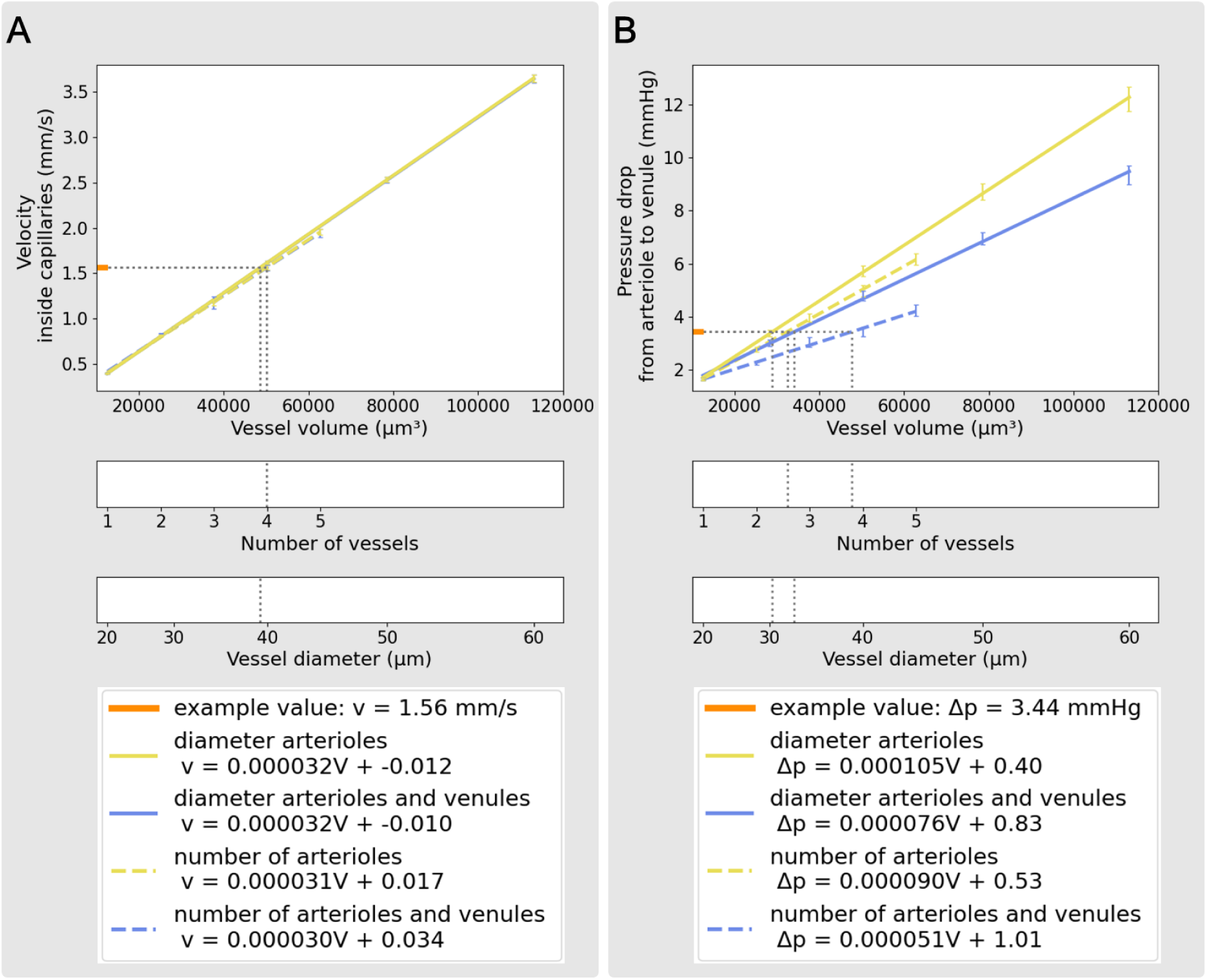
Regression analyses quantify the relationship between target values and the volume of arterioles and venules in our models. The example value pair corresponds to the simulation outcomes from a model with each four arterioles and venules, all with a diameter of 20 µm. The increase in vessel volume within the different model set corresponds to changes in either the number (dashed) or the diameter (continuous) of only arterioles (yellow) or both arterioles and venules (blue). (A) In the case of mean flow velocity within capillaries, the regression lines of the different model sets overlap. (B) The regression lines for pressure drop yield identifiable variations. All results are from simulations using the Newtonian viscosity model.

In summary, we found that the target values increase linearly with the arteriolar volume. With our current choice of boundary conditions, the simulation results are consistent with the existing literature values for flow velocity and pressure drop. Comparison of the latter with our results from Newtonian simulations suggests that the ACN of a single alveolus is linked to at least arterioles with diameters of 20 µm or a single arteriole with a minimum diameter of 30 µm. On the other hand, comparing our non-Newtonian simulation results with literature values indicate that the ACN of a single alveolus is connected to no more than one 20-µm-arteriole. If corresponding experimental measurements for capillary velocity and pressure drop were available, features of the structure of the connection between the ACN and the vascular tree could be inferred.

## 4 Discussion

By synergistically combining data-based 3D morphological modeling and computational fluid dynamics simulations, we developed an integrative approach to study the morphology and connectivity of the alveolar capillary network.

### Integrative 3D modeling and evaluation of alveolar capillary network geometry

With regard to the morphological aspect, 3D *in silico* modeling supports the concept of the sheet-flow model geometry as an appropriate representation of the ACN. We integrated data from several morphological studies (Hansen and Ampaya, 1975; Mercer et al., 1994; Ochs et al., 2004; Kawakami and Takizawa, 1987; Toshima et al., 2004; Matsuda et al., 1987; Stone et al., 1992; Mühlfeld et al., 2010; Gehr et al., 1978), using some parameters as a basis for 3D modeling and the others for model evaluation. The alveolar capillary network was constructed using two concepts, sheet-flow and tube-flow. Both model types yielded volume and surface area values below published estimates. Variations in the data such as these were anticipated. Experimental methods for studying alveolar morphology involve tissue fixation, which inherently introduce bias and loss of information (Weibel, 1984). Other uncertainties arise from simplifications like assuming alveoli to be 3/4 spheroids or from having to derive some reference values from other parameters. For example, the volume of the alveolar capillary bed was estimated from the capillary volume of the entire lung (Mühlfeld et al., 2010) and a number of 480 million alveoli in the lung (Ochs et al., 2004). Despite these limitations, the experimental data is extremely valuable and the only source on which we can draw to date. Nevertheless, we argue that the parameters characterizing alveolar morphology should not be considered in isolation. Instead, they should be related to each other, as we have done in our integrative 3D model. In this way, dependencies among the different parameters (e.g., between diameter or volume of the alveolus and surface area of the ACN) can be quantified and the conclusiveness of the measurements can be assessed.

### CFD simulation of blood flow in alveolar capillaries: model evaluation and viscosity impact

Adding CFD simulations to the sheet-flow geometry, we estimated blood flow paths, velocity and pressure distribution in three spatial dimensions.

We observed a linear correlation between the inlet velocity boundary condition and our target values - mean flow velocity within capillaries and pressure drop from arteriole to venule. In addition, the pressure drop was dependent on the fluid viscosity. Consequently, comparison between simulations with Newtonian and non-Newtonian viscosity models yielded comparable velocity values (around 0.4 mm/s) but considerable differences in pressure drop (1.64 mmHg with a Newtonian fluid versus 3.54 mmHg with a non-Newtonian fluid). In general, the results showed a decrease of the flow velocity inside the capillaries compared to inside the supply vessels and that the flow direction parallel to the inlet and outlet vessels was strongly preferred. Our simulation results are consistent with published *in vitro* measurements. Stauber et al. (2017) have developed a device that mimics the sheet-flow ACN. For 7 µm interpillar distance, the closest match to our sheet-flow model, they have measured a red blood cell (RBC) flow velocity of 0.3 mm/s at a pressure gradient of 0.5 kPa (3.75 mmHg). Previous theoretical work has predicted distribution of blood flow in line with our results (Zurita and Hurtado, 2022). Their 3D hollow sphere model compares well with our sheet-flow ACN models, except that the geometry is based on mouse morphometrics and is more simplified. They observed a decrease in flow velocity in the middle of the capillary bed and higher velocities near the inlet and outlet surfaces, as well as a continuous pressure drop from inlet to outlet. Burrowes et al. (2004) have predicted blood flow in 3D tube-flow ACN models across several alveoli, employing a fixed pressure drop of 8 cmH_2_O (5.88 mmHg) as a boundary condition. The mean flow velocity, derived from transit times, ranged from 0.15 mm/s to 1.07 mm/s for different transmural pressure conditions. In summary, the flow velocity within capillaries in our simulations lay within the range of references from previous theoretical and *in vitro* studies. Furthermore, the reference values for arteriole-to-venule pressure drop were undercut in our CFD simulations with constant viscosity but closely approximated in simulations with the non-Newtonian viscosity model. The selection of a 2 cP constant viscosity in our Newtonian CFD simulations was based on estimated apparent viscosities of blood in pulmonary capillaries from several studies (Fung, 1984; Kiani and Hudetz, 1991; Fung, 1997; Grotberg and Romanò, 2023). In selecting the non-Newtonian viscosity model and the corresponding parameters, we followed the approach of Albors et al. (2023) who used this model to simulate blood flow in the left atrium. However, it covers a range of viscosity values between 3.5 cP and 56 cP which is above the estimated apparent viscosity in the pulmonary capillaries. In the future, it is advisable to use a viscosity model that accurately represents blood’s rheological properties within the microvasculature, where factors like hematocrit and vessel diameter exert a strong influence.

To determine the most suitable model assumptions, including but not limited to the boundary conditions and the viscosity model, we rely on a consistent data set, ideally derived from the same experiment or, at a minimum, from the same species. Currently, literature values for the choice of boundary conditions and as benchmarks for comparison with the simulation results have to be taken from several studies working with different species. These physiological metrics sometimes show large differences between different species. This is complicated by the fact that these measurements were performed in vessels of varying diameters. The optimal data set would include blood flow velocities in arterioles and capillaries, blood pressure in arterioles and venules, and the mean diameters of the vessels in which these values were measured.

In summary, we have developed a base model for simulating blood flow in the pulmonary capillaries. Qualitatively, our simulation results are consistent with those from previously published *in vitro* studies and blood flow simulations in the literature. However, we observed strong dependencies of our target values on the inlet velocity boundary condition and / or the fluid viscosity. In order to refine and validate our model, appropriate experimental data would be needed.

### Investigating ACN connectivity with the vascular tree: A novel approach linking morphology and function

For connectivity analyses, we employed our integrative *in silico* approach to investigate the connection of the ACN to arterioles and venules. The CFD results for mean flow velocity inside the capillary bed and the pressure drop from arteriole to venule in our connectivity models were within the range of published reference values.

Comparison of our simulation results on flow velocity with literature references suggests that a single alveolar ACN is likely connected to at least two arterioles with diameters of 20 µm or a single arteriole with a diameter of at least 30 µm. The pressure drop results depended on the viscosity model. With constant low viscosity (Newtonian model), a similar prediction can be made about the arteriole configuration as for velocity simulation results. In contrast, simulations with the non-Newtonian model indicate that the ACN around an alveolus is connected to no more than one 20-µm-arteriole.

Previous studies that have examined the linkage between arterioles, venules, and the ACN in the human lung have done so from a morphological perspective (Huang et al., 1996; Pump, 1966; Horsfield, 1978). Across these studies, naming of the smallest pulmonary vessels is inconsistent, but considering vessel diameter allows for direct comparison. These findings suggest that one arteriole with a diameter of approximately 20 µm connects to multiple alveoli, with values ranging from 4.8 (Huang et al., 1996) to 24 (Horsfield, 1978). For the actual connection between the vascular tree and the capillary network so-called precapillary vessels have been introduced (Horsfield, 1978; Pump, 1966). Only vague information is provided about these vessels, such as that their diameters approach the capillary diameter.

The morphological estimates suggest that in our simulations we should observe an oversupply of the capillary network around a single alveolus already by a single 20 µm arteriole. However, our results do not indicate this. Once experimental data becomes available to further verify our CFD models, it will be intriguing to examine whether the predictions from morphological studies still hold from a functional point of view.

Our current model shows possibilities for predicting morphological features based on physiological parameters. If consistent experimental measurements for flow velocity and pressure drop were available, details of the arteriole and venule configuration could be inferred. Depending on the values, we can identify two possible connectivity models based on either the number or the diameter of arterioles or we can make a unique prediction.

Previous theoretical studies on pulmonary blood flow focused on predicting physiological parameters based on morphological properties. The work of Zurita and Hurtado (2022) has predicted the 3D spatial distribution of blood flow and gas exchange. Multi-scale models (Clark et al., 2010, 2011; Clark and Tawhai, 2018; Ebrahimi et al., 2022) are more complex and allow to investigate even broader relationships. They enable predictions of both whole lung function with alterations in small vascular structures and small-scale vascular function in response to impairment of the proximal pulmonary artery. Theoretical studies with CFD blood flow simulations for other organs such as the heart are even more elaborated (Totorean et al., 2022). Mill et al. (2021b,a); Pons et al. (2022) have employed CFD simulations with patient-specific boundary conditions and dynamic mesh motion of the 3D geometry to assess thrombus risk in the left atrial appendage, showcasing the clinical potential of CFD in hemodynamic studies.

The novelty in our approach is that we exploit the interdependence of structure and function in the opposite direction: On the basis of physiological blood flow parameters, we evaluated the realism of systematically introduced morphological variations. To our knowledge, such an inverse approach on structure-function- relationships has only ever been employed on a cellular level, inferring position and morphology of neurons from *in vivo* extracellular voltage recordings using biophysical modeling (Chen et al., 2021, 2023).

### Conclusion and outlook

In conclusion, our integrative *in silico* modeling approach complements previous studies and offers a unique perspective on the structure-function relationship of the pulmonary microvasculature. By synergistically combining data-based 3D modeling and computational fluid dynamics simulations, we successfully refined our understanding of the alveolar capillary network and its relationship with the vascular tree. Moving forward with appropriate experimental data for validation and incorporating a suitable viscosity model to account for blood’s rheological behavior, our *in silico* model holds promise for more precise simulations and further insights into microvascular hemodynamics. In addition, we plan to incorporate these findings into our interactive platform for simulating gas exchange in the human alveolus (Schmid et al., 2022). For future research, introducing dynamic changes in alveolar volume over time will add complexity to the model, enhancing its fidelity to real physiological scenarios. The same holds true for the combination of several alveoli and their mechanical as well as hemodynamic interdependence. Furthermore, automating the morphological modeling process would facilitate more efficient exploration of systematic geometric variation.

On a final note, our approach demonstrates the feasibility of deriving morphological aspects from physiological parameters in the case of blood flow simulations in the pulmonary microvasculature. A new strategy that can be transferred to other tissues and organs.

## Acknowledgements

K.S. was supported by an EMBO Scientific Exchange Grant (9271). K.S. and S.C.F. acknowledge the support by a grant from Universitätsbund Würzburg (AZ21-16). A.C.H. was supported by German Research Foundation (DFG, SFB-TR 84, B6, Z1a) as well as by the Einstein Foundation (EC3R) Charité 3R.

## Supplementary Material

### S.1 3D modeling of the alveolar capillary network in Blender

In Blender, different workflows were used to create the tube-flow and sheet-flow models of the ACN (Figure S.1). As a first step, a surface was extended around the alveolar base, serving as the foundation for the later capillary network. In the case of tube-flow modeling, this surface was positioned at the distance of the capillary radius from the alveolar base (A-I in Figure S.1). In published electron microscopy images of the ACN (Mühlfeld, 2021; Du et al., 2020; Buchacker et al., 2019), we observe mainly trivalent nodes between capillary segments. Therefore, we chose a topology of hexagonal and pentagonal faces for the capillary base surface, resulting in trivalent vertices. We obtained this topology using the Dual Mesh algorithm of the Tissue add-on in Blender. This algorithm generates a polygonal mesh from a triangular mesh by subdividing the faces and then dissolving the original vertices. The edges of the faces would form the centerlines of the later capillary tubes. This would result in voids in the center of the faces, representing the capillary loops or interalveolar pores of Kohn, depending on their diameter. When modeling sheet-flow, the capillary base surface was directly at the alveolar base (B-I in Figure S.1). Positions for tissue pillars were randomly distributed on this surface using a simple script so that, on average, they were at the target distance from each other. The minimum distance was one and a half times the pillar diameter. In addition, positions for a required number of pores of Kohn were manually distributed. At these positions, the capillary base surface was perforated with holes corresponding to the diameters of tissue pillars and pores of Kohn, respectively. Subsequently, the capillary base surface was transformed into a voluminous body. In the tube-flow model, this was achieved using the Wireframe modifier, which iterates over all faces and converts edges to four-sided polygons (A-II in Figure S.1). The width of these polygons, specified by the Thickness setting of the modifier, determines the capillary diameter. For the sheet-flow model, the capillary base surface was extruded away from the alveolar base by the desired sheet thickness (B-II in Figure S.1). Finally, the capillary geometry was smoothed by two iterations of the Subdivision Surface modifier (Figure S.1 step III).

**Figure S.1:**
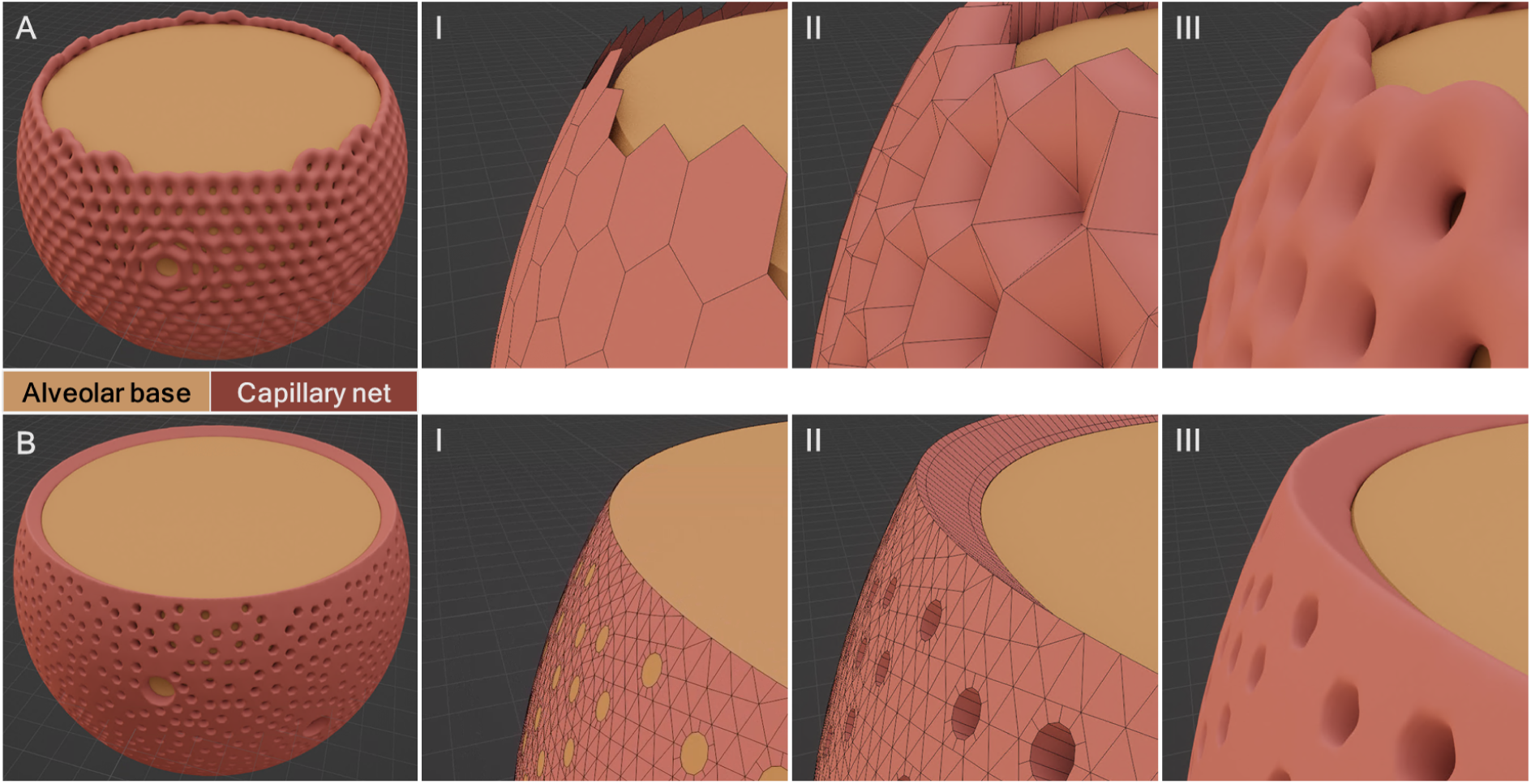
Different modeling workflows in Blender were used to create the tube-flow (A) and sheet-flow (B) models of the alveolar capillary network. Step I. A surface was extended around the alveolar base (brown), serving as the foundation for the later capillary network (red). Step II. The capillary base surface was transformed into a voluminous body. Step III. The capillary geometry was smoothed by two iterations of the “subdivision surface” modifier.

### S.2 Volumetric meshing with Ansys Fluent Meshing

The workflow in Ansys Fluent Meshing consisted of three main steps, in which the surface mesh, the boundary layer and the volumetric mesh were built up one after the other. The surface mesh was defined along the capillary wall and inlet and outlet surfaces based on a curvature size function. This method controls the distribution of mesh size on a surface based on the following user-defined parameters: Minimum and maximum values for the size of the mesh edges, a “growth rate” defining the maximum size difference between adjacent mesh elements, and the curvature normal angle. The curvature normal angle defines the angle of curvature of the surface that can be covered by a single element. The result was a surface mesh with a particularly high resolution in regions of curvature. In a next step, the boundary layer of prism elements was generated starting from the surface mesh at the wall zone. Here, the “last ratio” meshing method was chosen. This method controls the distribution of the prism layers depending on the last, innermost layer, which forms the transition to the polyhedral elements. For this last prism layer, the ratio of height to length of the individual elements, the so-called transition ratio, is determined. In addition, the number of layers and the height of the first layer (closest to the wall) was specified. Starting from this, the prism layers were distributed so that their height increases continuously from the wall to the center of the volume. Finally, the remaining volume was filled with polyhedra. The size function settings for the volumetric mesh, i.e., element size maximum and growth rate, were chosen as for the surface mesh. Two FV meshes of different complexity were created for each ACN model geometry. The input parameters for the meshing workflow were adjusted from the coarse mesh to the fine mesh using a scaling factor of 1.6 (Table S.1).

**Table S.1:** FV meshes for computational fluid dynamics were created with the Watertight Geometry workflow in Ansys Fluent Meshing. Two meshes of different complexity were generated by scaling the input parameters with a refinement factor 1.6.

|  | Coarse mesh | Fine mesh |
| --- | --- | --- |
| **Surface mesh** |  |  |
| Minimum size ($\mu\text{m}$) | 0.02 | 0.0125 |
| Maximum size ($\mu\text{m}$) | 2 | 1.25 |
| Growth rate | 1.2 | 1.2 |
| Curvature normal angle ($^\circ$) | 9.6 | 6 |
| **Boundary layer** |  |  |
| Number of layers | 6 | 10 |
| Transition ratio | 0.5 | 0.5 |
| First height ($\mu\text{m}$) | 0.02 | 0.0125 |
| **Volumetric mesh** |  |  |
| Growth rate | 1.2 | 1.2 |
| Max cell length ($\mu\text{m}$) | 2 | 1.25 |

### S.3 Finite volume mesh quality

FV mesh quality was evaluated based on three different parameters (Table S.2). The first was inverse orthogonal quality (IOQ). It quantifies how far a mesh cell deviates from orthogonality with respect to its faces. It can have values between 0 (ideal) and 1 (poor) where a value of 0 would mean that all cell angles are 90°. The IOQ of a mesh has a direct influence on the discretization error and iterative convergence of numerical solutions, as well as on the scalability of the mesh. All elements of the mesh had to comply with their IOQ below the limit of 0.95. Their mean IOQ ranged from 0.16 to 0.19. The second parameter was size change, i.e., the increase in volume between neighboring cells. This value should be kept as low as possible, ideally close to 1, to avoid errors in interpolation between control volumes. The mean size change in our FV meshes was between 3.8 and 3.9. Lastly, the aspect ratio was considered. This parameter is a measure of the ratio of the length and height of the cells. It does not have a strong effect on the discretization error, but on rounding errors. The mean aspect ratio of the meshes used in this work ranged from 18 to 26.

**Table S.2:** The quality of finite volume meshes was evaluated based on the parameters: inverse orthogonal quality (IOQ), size change, and aspect ratio.

| Mesh size and quality | Coarse mesh | Fine mesh |
| --- | --- | --- |
| Number of cells ($10^6$) | 4.9 | 7.6 |
| IOQ | 0.17 | 0.2 |
| Size change | 3.88 | 3.92 |
| Aspect ratio | 18.0 | 26.4 |

### S.4 CFD simulation setup in Ansys

**Table S.3:** Parameter settings for the computational fluid dynamics simulations in Ansys^®^ Academic Research CFD, release 2022 R2.

|  |  |
| --- | --- |
| **General** |  |
| solver type | pressure-based |
| time | steady |
| viscous model | laminar |
| **Material Settings** |  |
| type | fluid |
| density | 1050 kg/m<sup>3</sup> |
| viscosity | 0.002 kg/(m·s) or Carreau model (Equation 3) |
| **Boundary Conditions** |  |
| inlet | velocity magnitude 0.0023 m/s |
| wall | stationary, no slip |
| outlet | gauge pressure 1127 Pa |
| **Solution Methods** |  |
| pressure-velocity coupling - scheme | coupled |
| spatial discretization - gradient | green-gauss node based |
| spatial discretization - pressure | second order |
| spatial discretization - momentum | second order upwind |

### S.5 CFD quality assurance

CFD is a numerical approach, and its results always represent an approximation to the unknown exact solution. To ensure the highest quality in CFD simulations, it is important to assess the accuracy of this approximation. This involves identifying errors in the physical model, which requires a comparison with suitable experimental data. Before this, however, it is essential to monitored and mitigated the following systematic errors: Computer rounding error, iterative convergence error and discretization error.

#### Computer rounding error

The computer rounding error depends on the accuracy with which floating numbers are stored on the computer. Compared to the other errors, it is usually not significant. In our true-to-scale models, we calculate with double precision, as recommended by Ansys for FV meshes with very small minimum face areas. (The minimum face area of our meshes is in the scale of 10^-19^ m^2^.)

#### Iterative convergence error

The iterative convergence error is defined as the difference between the numerical and the exact solution of the discretized flow problem and its magnitude is contingent upon the point at which the iteration process is concluded. In order to identify an optimal termination point that minimizes the iteration error, specific sensors are monitored and convergence criteria are introduced. These sensors comprise residual values, solution imbalances and coefficients of variation of target values (Figure S.2).

The residual measures the local imbalance of a variable in each control volume. For every model parameter, the residual values for all control volumes are summed, scaled by a scaling factor representative of the flow rate of this variable, and normalized. The iteration process is continued until the residual norms fall below a specific threshold. The other sensors were checked according to the “human-in-the-loop” principle. Once a simulation had converged, we manually verified if the global mass balance had leveled out around zero (values were in the range of 10^-18^ to 10^-13^ kg/s) and if our target values had stabilized at a constant value (coefficients of variations ranging from 8.2*·*10^-8^ to 1.5*·*10^-4^ for the mean capillary velocity and from 7.6*·*10^-7^ to 3.1*·*10^-4^ for the pressure drop).

#### Discretization error

The discretization error is defined as the difference between the solution on a mesh with a certain width (size of the control volumes) and the solution on an infinitely fine mesh. It approaches its minimum as the mesh width decreases and the number of mesh elements, i.e., the mesh complexity, increases. In order to estimate the discretization error of the CFD solutions, two meshes with distinct complexity were compared (Table S.1). The CFD results presented in this work were obtained with the fine meshes. The discretization error e*_h_* of the solution on a fine mesh (f*_fine_*) was estimated using the solution on the corresponding coarse mesh f*_coarse_* as follows:

**Figure S.2:**
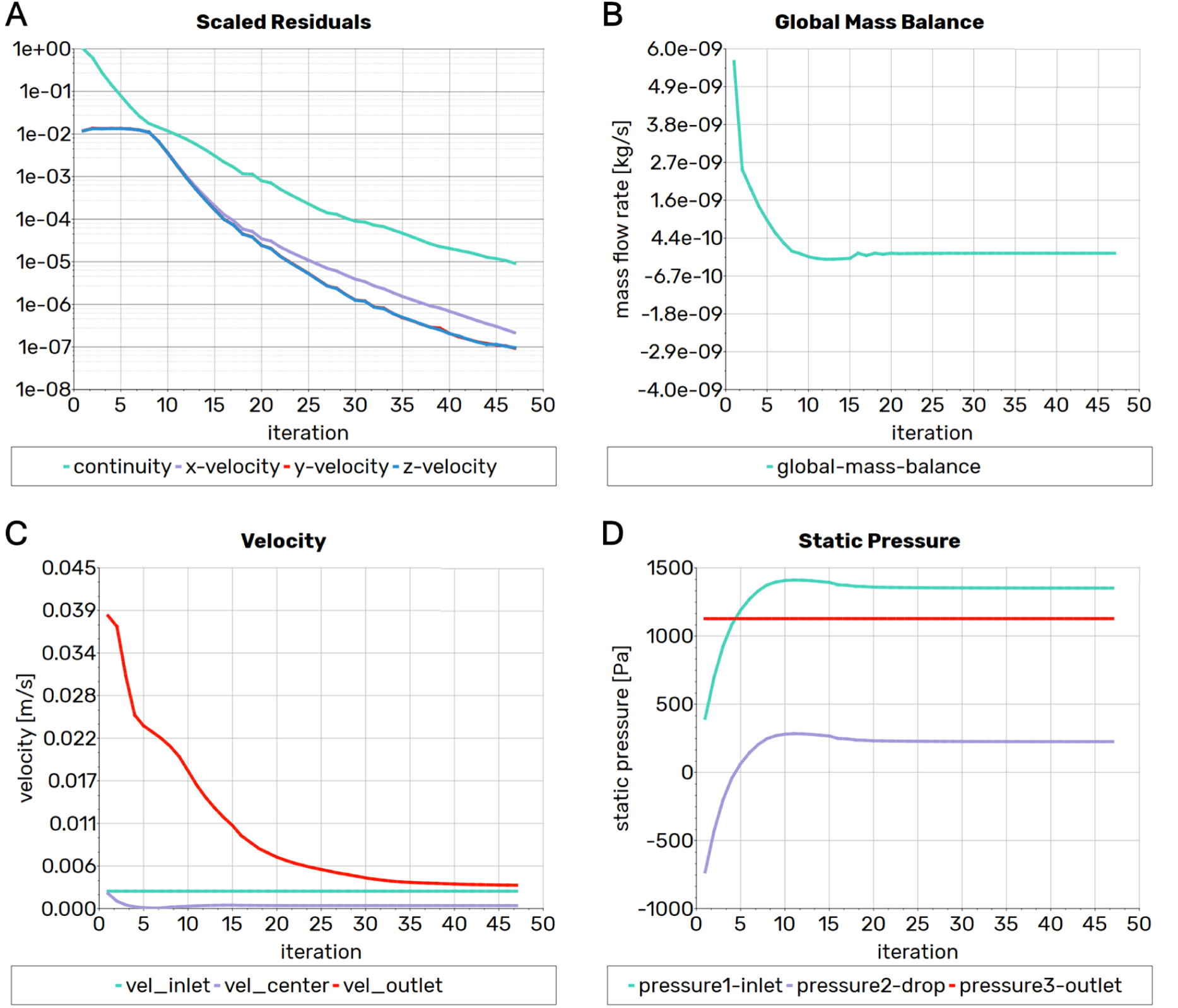
The convergence of the computational fluid dynamics solution was monitored with several sensors. Residual values of model variables (A) measure their imbalance in the finite volume equations. A convergence criterion was that these residuals (scaled and normalized) fall below a threshold of 1*·*10^-5^. Solution imbalances were observed with the global mass balance (B), defined as the sum of mass flow rates of all inlet and outlet faces. In this example case, the global mass balance was -1.3*·*10^-15^ kg/s when the iteration process was terminated. In addition, it was checked whether the target values mean flow velocity (C) and pressure drop had (D) also converged. In this example case, the mean flow velocity had stabilized at 0.398 mm/s at the end of the simulation. The coefficient of variation over the last ten iterations was 5.7*·*10^-5^. The pressure drop was 1.68 mmHg (224 Pa) with a coefficient of variation of 3.1*·*10^-5^ over the last ten iterations.

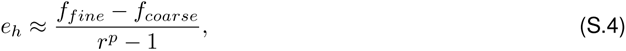

where r is the refinement factor between the two meshes and p is the order of the discretization scheme.

In our simulations, the discretization error lay within the range of 0.15 % to 0.39 % of the result value for the mean velocity within the capillaries and from 0.09 % to 0.95 % of the result value for the pressure drop between arteriole and venule.

### S.6 Connectivity analyses

**Figure S.3:**
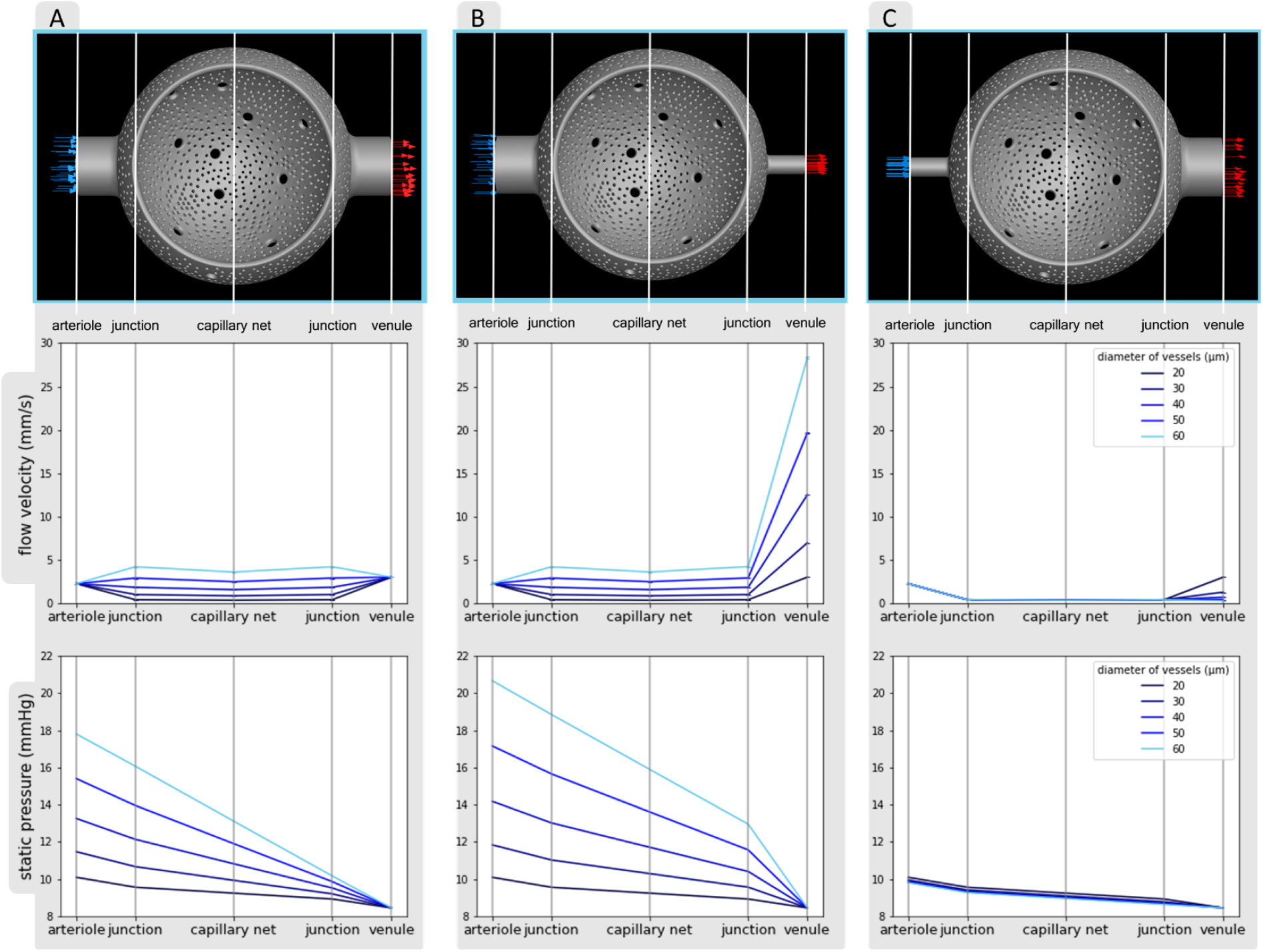
Flow velocity and static pressure were measured at different positions: At the arteriole (inlet), at the junction between arteriole and capillary net, at the center of the capillary net, at the junction between capillary net and venule, and at the venule (outlet). The results were compared between models with varying diameters of both arterioles and venules (A), only arterioles (B), or only venules (C). 3D models with 60 µm diameter vessels (top row) are shown as examples, the positions of the measurements are indicated by white lines.

**Figure S.4:**
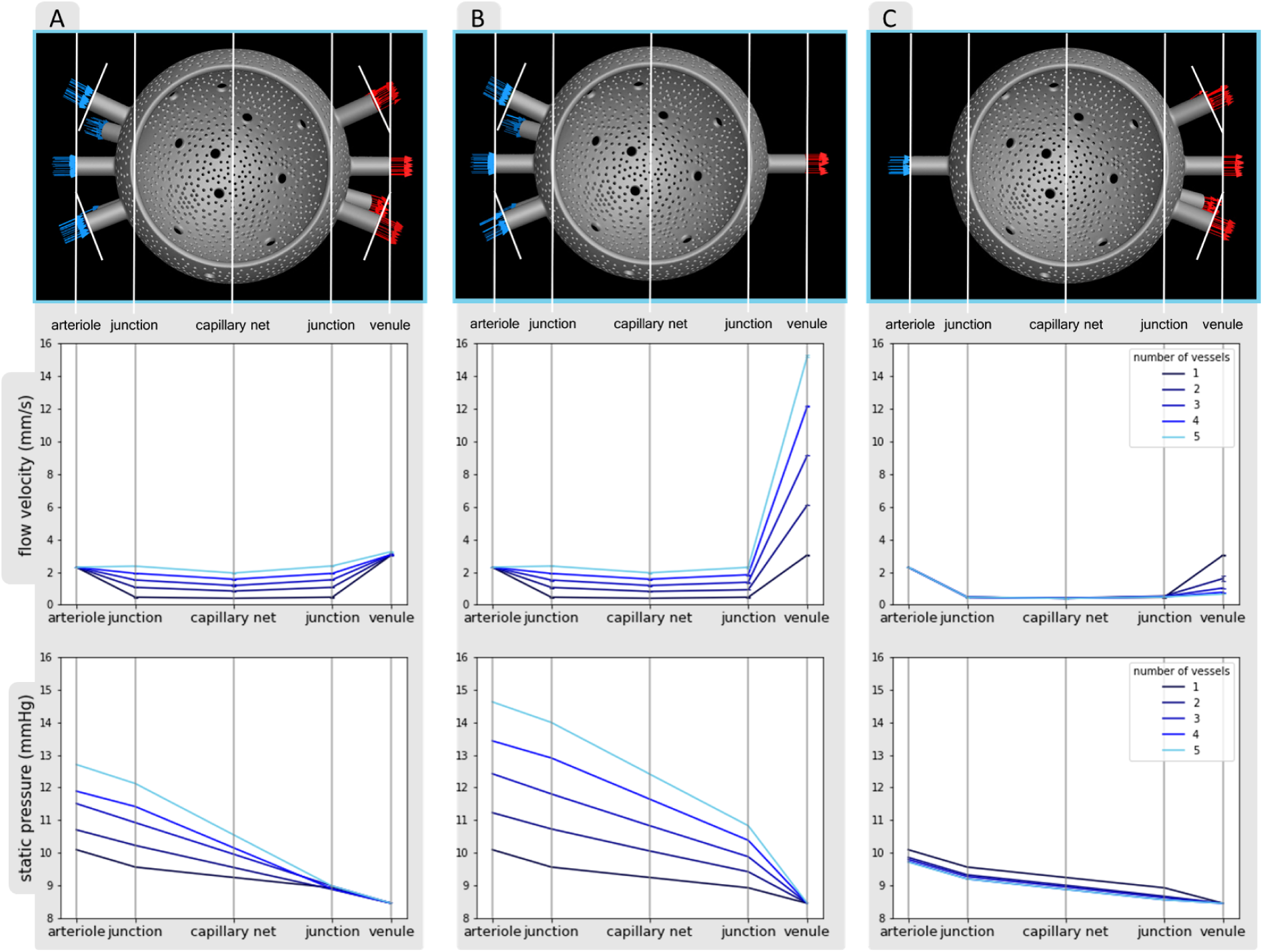
Flow velocity and static pressure were measured at different positions: At the arteriole (inlet), at the junction between arteriole and capillary net, at the center of the capillary net, at the junction between capillary net and venule, and at the venule (outlet). The results were compared between models with varying numbers of both arterioles and venules (A), only arterioles (B), or only venules (C). 3D models with five vessels (top row) are shown as examples, the positions of the measurements are indicated by white lines.

**Table S.4:** Hemodynamic indices from computational fluid dynamics simulations using a Newtonian viscosity model in models with different configurations of the number of arterioles and venules. Mean values and standard deviations are given for flow velocity within capillaries and pressure drop from arteriole to venule.

|  | Vessel number |  |  |  |  |
| --- | --- | --- | --- | --- | --- |
|  | 1 | 2 | 3 | 4 | 5 |
| **Mean flow velocity within capillaries (mm/s) (±SD)** |  |  |  |  |  |
| Arterioles and Venules | 0.39 (±0.004) | 0.84 (±0.007) | 1.18 (±0.07) | 1.56 (±0.03) | 1.94 (±0.04) |
| Only Arterioles | 0.39 (±0.004) | 0.81 (±0.009) | 1.18 (±0.04) | 1.57 (±0.03) | 1.96 (±0.03) |
| Only Venules | 0.39 (±0.004) | 0.41 (±0.004) | 0.39 (±0.01) | 0.39 (±0.005) | 0.39 (±0.006) |
| **Mean arteriole-to-venule pressure drop (mmHg) (±SD)** |  |  |  |  |  |
| Arterioles and Venules | 1.64 (±0.04) | 2.25 (±0.07) | 3.06 (±0.18) | 3.44 (±0.16) | 4.25 (±0.21) |
| Only Arterioles | 1.64 (±0.04) | 2.78 (±0.10) | 3.97 (±0.15) | 4.98 (±0.21) | 6.17 (±0.21) |
| Only Venules | 1.64 (±0.04) | 1.41 (±0.05) | 1.34 (±0.05) | 1.28 (±0.05) | 1.26 (±0.05) |

**Table S.5:** Hemodynamic indices from computational fluid dynamics simulations using a Newtonian viscosity model in models with different configurations of the diameter of arterioles and venules. Mean values and standard deviations are given for flow velocity within capillaries and pressure drop from arteriole to venule.

| | Vessel diameter (μm) | | | | |
| --- | --- | --- | --- | --- | --- |
|  | 20 | 30 | 40 | 50 | 60 |
| **Mean flow velocity within capillaries (mm/s) (±SD)** |  |  |  |  |  |
| Arterioles and Venules | 0.39 (±0.004) | 0.91 (±0.01) | 1.61 (±0.02) | 2.53 (±0.03) | 3.64 (±0.04) |
| Only Arterioles | 0.39 (±0.004) | 0.91 (±0.01) | 1.62 (±0.02) | 2.53 (±0.03) | 3.65 (±0.04) |
| Only Venules | 0.39 (±0.004) | 0.39 (±0.004) | 0.39 (±0.004) | 0.39 (±0.004) | 0.39 (±0.004) |
| **Mean arteriole-to-venule pressure drop (mmHg) (±SD)** |  |  |  |  |  |
| Arterioles and Venules | 1.64 (±0.04) | 3.02 (±0.12) | 4.81 (±0.18) | 6.95 (±0.24) | 9.35 (±0.35) |
| Only Arterioles | 1.64 (±0.04) | 3.39 (±0.12) | 5.73 (±0.19) | 8.70 (±0.31) | 12.21 (±0.46) |
| Only Venules | 1.64 (±0.04) | 1.50 (±0.06) | 1.43 (±0.04) | 1.39 (±0.05) | 1.35 (±0.05) |

**Figure S.5:**
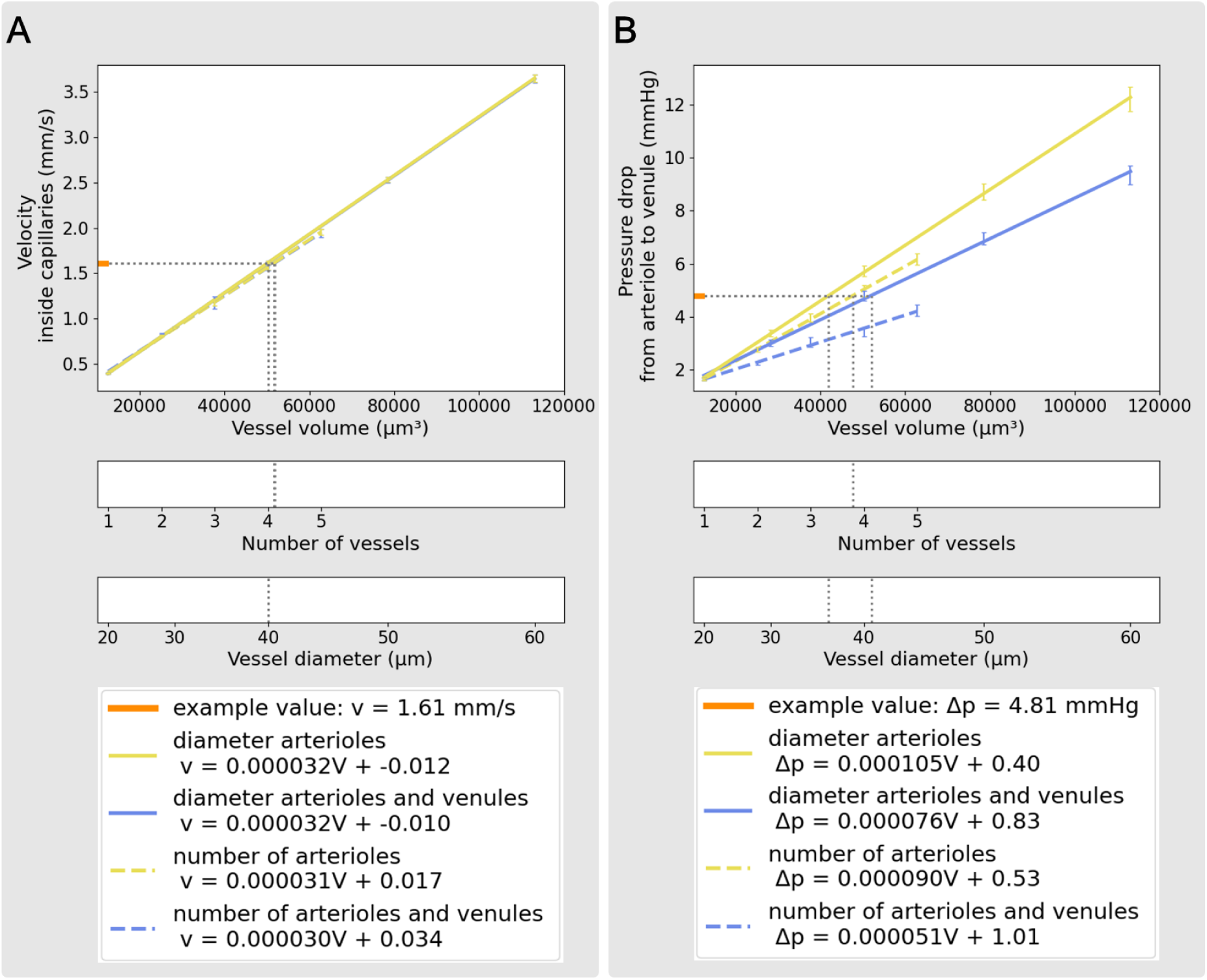
Regression analyses quantify the relationship between target values and the volume of arterioles and venules in our models. The example value pair corresponds to the simulation outcomes from a model with one arteriole and one venule, both with a diameter of 40 µm. The increase in vessel volume within the different model sets corresponds to changes in either the number (dashed) or the diameter (continuous) of only arterioles (yellow) or both arterioles and venules (blue). (A) In the case of mean flow velocity within capillaries, the regression lines of the different model sets overlap. (B) The regression lines for pressure drop yield identifiable variations. All results are from simulations using the Newtonian viscosity model.

### S.7 CFD simulations with non-Newtonian viscosity model

**Figure S.6:**
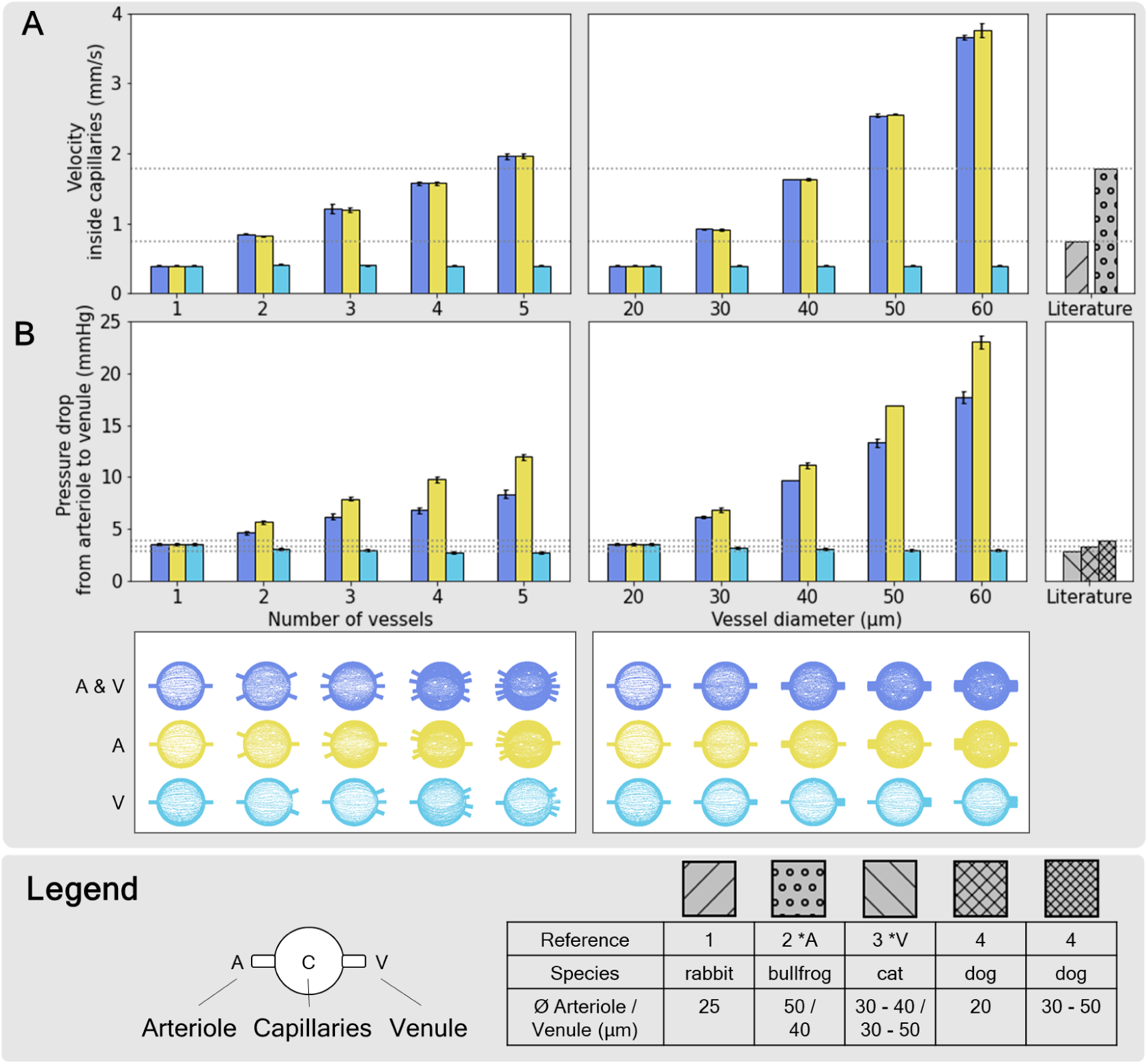
*In silico* hemodynamic indices from flow simulations in symmetric and asymmetric connectivity models. Alveolar capillary network models were connected to increasing numbers (left panels) or diameters (right panels) of arterioles (yellow), venules (turquoise), or both (blue). Flow velocity within capillaries (A) and pressure drop from arteriole to venule (B) were predicted with computational fluid dynamics simulations using a non-Newtonian viscosity model. The results were compared with literature values of ^1^rabbit (Schlosser et al., 1965), ^2^bullfrog (Horimoto et al., 1979), ^3^cat (Nagasaka et al., 1984) and ^4^dog (Bhattacharya and Staub, 1980). **\***A Simulation boundary condition for inlet velocity at the arterioles were based on a reference from this publication. **\***V Simulation boundary condition for outlet pressure at the venules was based on references from this publication.

## Notes

### Competing Interest Statement

The authors have declared no competing interest.

## References

Carlos Albors, Andy L. Olivares, Xavier Iriart, Hubert Cochet, Jordi Mill, and Oscar Camara. Impact of Blood Rheological Strategies on the Optimization of Patient-Specific LAAO Configurations for Thrombus Assessment. In Olivier Bernard, Patrick Clarysse, Nicolas Duchateau, Jacques Ohayon, and Magalie Viallon, editors, Functional Imaging and Modeling of the Heart, Lecture Notes in Computer Science, pages 485–494, Cham, 2023. Springer Nature Switzerland. ISBN 978-3-031-35302-4.

Inc. ANSYS. Continuity and Momentum Equations. In Ansys® Fluent Theory Guide, Help System. Release 12.0 edition, January 2009.

Inc. ANSYS. Ansys® Academic Research CFD, 2022.

J. Bhattacharya and N. C. Staub. Direct Measurement of Microvascular Pressures in the Isolated Perfused Dog Lung. Science, 210(4467):327–328, October 1980. doi: 10.1126/science.7423192. Publisher: American Association for the Advancement of Science.

R. J. Bosman, Ch. P. Stoutenbeek, and D. F. Zandstra. Non-invasive pulmonary blood flow measurement by means of CO2 analysis of expiratory gases. Intensive Care Medicine, 17(2):98–102, February 1991. ISSN 1432-1238. doi: 10.1007/BF01691431. URL 10.1007/BF01691431.

S. Braber, K. a. T. Verheijden, P. a. J. Henricks, A. D. Kraneveld, and G. Folkerts. A comparison of fixation methods on lung morphology in a murine model of emphysema. American Journal of Physiology-Lung Cellular and Molecular Physiology, 299(6):L843–L851, December 2010. ISSN 1040-0605. doi: 10.1152/ ajplung.00192.2010. Publisher: American Physiological Society.

Tobias Buchacker, Christian Mühlfeld, Christoph Wrede, Willi L. Wagner, Richard Beare, Matt McCormick, and Roman Grothausmann. Assessment of the Alveolar Capillary Network in the Postnatal Mouse Lung in 3D Using Serial Block-Face Scanning Electron Microscopy. Frontiers in Physiology, 10, 2019. ISSN 1664-042X. URL https://www.frontiersin.org/articles/10.3389/fphys.2019.01357.

Kelly S. Burrowes, Merryn H. Tawhai, and Peter J. Hunter. Modeling RBC and Neutrophil Distribution Through an Anatomically Based Pulmonary Capillary Network. Annals of Biomedical Engineering, 32(4):585–595, April 2004. ISSN 1573-9686. doi: 10.1023/B:ABME.0000019178.95185.ad.

Ziao Chen, Dan Dopp, Satish S Nair, and Drew B Headley. Inferring Morphology of a Neuron from In Vivo LFP Data. In 2021 10th International IEEE/EMBS Conference on Neural Engineering (NER), pages 774–777, May 2021. doi: 10.1109/NER49283.2021.9441161. URL https://ieeexplore.ieee.org/abstract/document/9441161. ISSN: 1948-3554.

Ziao Chen, Matthew Carroll, and Satish S. Nair. Inferring Pyramidal Neuron Morphology using EAP Data. International IEEE/EMBS Conference on Neural Engineering: [proceedings]. International IEEE EMBS Conference on Neural Engineering, 2023, April 2023. ISSN 1948-3546. doi: 10.1109/ner52421.2023. 10123903.

A. R. Clark and M. H. Tawhai. TEMPORAL AND SPATIAL HETEROGENEITY IN PULMONARY PERFUSION: A MATHEMATICAL MODEL TO PREDICT INTERACTIONS BETWEEN MACRO- AND MICRO-VESSELS IN HEALTH AND DISEASE. The ANZIAM Journal, 59(4):562–580, April 2018. ISSN 1446-1811, 1446-8735. doi: 10.1017/S1446181118000111. Publisher: Cambridge University Press.

A. R. Clark, K. S. Burrowes, and M. H. Tawhai. Contribution of serial and parallel microperfusion to spatial variability in pulmonary inter- and intra-acinar blood flow. Journal of Applied Physiology, 108(5):1116–1126, May 2010. ISSN 8750-7587. doi: 10.1152/japplphysiol.01177.2009. Publisher: American Physiological Society.

A. R. Clark, M. H. Tawhai, E. A. Hoffman, and K. S. Burrowes. The interdependent contributions of gravitational and structural features to perfusion distribution in a multiscale model of the pulmonary circulation. Journal of Applied Physiology, 110(4):943–955, April 2011. ISSN 8750-7587. doi: 10.1152/japplphysiol.00775.2010. Publisher: American Physiological Society.

Blender Online Community. Blender - a 3D modelling and rendering package, 2018. URL http://www.blender.org.

Amit Dhadwal, Barry Wiggs, Claire M. Doerschuk, and Roger D. Kamm. Effects of anatomic variability on blood flow and pressure gradients in the pulmonary capillaries. Journal of Applied Physiology, 83(5): 1711–1720, November 1997. ISSN 8750-7587. doi: 10.1152/jappl.1997.83.5.1711. Publisher: American Physiological Society.

Xiaohua Du, Xia Liu, and James Blackar Mawolo. The Architecture of Alveolar Capillaries in the Lungs of Bactrian Camels (Camelus bactrianus). International Journal of Morphology, 38(6):1779–1785, December 2020. ISSN 0717-9502.

Behdad Shaarbaf Ebrahimi, Haribalan Kumar, Merryn H. Tawhai, Kelly S. Burrowes, Eric A. Hoffman, and Alys R. Clark. Simulating Multi-Scale Pulmonary Vascular Function by Coupling Computational Fluid Dynamics With an Anatomic Network Model. Frontiers in Network Physiology, 2:867551, April 2022. ISSN 2674-0109. doi: 10.3389/fnetp.2022.867551.

M. Folke, L. Cernerud, M. Ekström, and B. Hök. Critical review of non-invasive respiratory monitoring in medical care. Medical and Biological Engineering and Computing, 41(4):377–383, July 2003. ISSN 1741-0444. doi: 10.1007/BF02348078. URL 10.1007/BF02348078.

(Version 0.20.1) FreeCAD. [Software], 2022. Retrieved from https://www.freecad.org/.

Y. C. Fung. Microcirculation. In Y. C. Fung, editor, Biodynamics: Circulation, pages 224–289. Springer, New York, NY, 1984. ISBN 978-1-4757-3884-1. doi: 10.1007/978-1-4757-3884-1.

Y. C. Fung. Circulation. In Biomechanics, pages XVIII, 572. Springer New York, 2 edition, 1997.

Y C Fung and S S Sobin. Theory of sheet flow in lung alveoli. Journal of Applied Physiology, 26(4):472– 488, April 1969. ISSN 8750-7587. doi: 10.1152/jappl.1969.26.4.472. Publisher: American Physiological Society.

Peter Gehr, Marianne Bachofen, and Ewald R. Weibel. The normal human lung: ultrastructure and morphometric estimation of diffusion capacity. Respiration Physiology, 32(2):121–140, February 1978. ISSN 0034-5687. doi: 10.1016/0034-5687(78)90104-4.

James B. Grotberg and Francesco Romanò. Computational pulmonary edema: A microvascular model of alveolar capillary and interstitial flow. APL Bioengineering, 7(3):036101, July 2023. ISSN 2473-2877. doi: 10.1063/5.0158324.

J. E. Hansen and E. P. Ampaya. Human air space shapes, sizes, areas, and volumes. Journal of Applied Physiology, 38(6):990–995, June 1975. ISSN 8750-7587. doi: 10.1152/jappl.1975.38.6.990.

Roland Hausmann, Horst Bock, Teresa Biermann, and Peter Betz. Influence of lung fixation technique on the state of alveolar expansion—a histomorphometrical study. Legal Medicine, 6(1):61–65, March 2004. ISSN 13446223. doi: 10.1016/j.legalmed.2003.08.007.

M. Horimoto, T. Koyama, H. Mishina, T. Asakura, and M. Murao. Blood flow velocity in pulmonary microvessels of bullfrog. Respiration Physiology, 37(1):45–59, May 1979. ISSN 0034-5687. doi: 10.1016/0034-5687(79)90091-4.

K Horsfield. Morphometry of the small pulmonary arteries in man. Circulation Research, 42(5):593–597, May 1978. doi: 10.1161/01.RES.42.5.593. Publisher: American Heart Association.

K. Horsfield and W. I. Gordon. Morphometry of pulmonary veins in man. Lung, 159(1):211–218, December 1981. ISSN 1432-1750. doi: 10.1007/BF02713917. URL 10.1007/BF02713917.

W. Huang, R. T. Yen, M. McLaurine, and G. Bledsoe. Morphometry of the human pulmonary vasculature. Journal of Applied Physiology, 81(5):2123–2133, November 1996. ISSN 8750-7587. doi: 10.1152/jappl. 1996.81.5.2123. Publisher: American Physiological Society.

Yaqi Huang, Claire M. Doerschuk, and Roger D. Kamm. Computational modeling of RBC and neutrophil transit through the pulmonary capillaries. Journal of Applied Physiology, 90(2):545–564, February 2001. ISSN 8750-7587. doi: 10.1152/jappl.2001.90.2.545. Publisher: American Physiological Society.

Autodesk Inc. Autodesk Meshmixer, 2019. URL http://www.meshmixer.com.

A. P. Javed, W. F. Whimster, M. H. Deverell, and M. J. Cookson. Distribution of alveolar wall per unit volume in the human lung. Analytical cellular pathology, 6(2):129–136, February 1994. ISSN 1878-3651.

M. Kawakami and T. Takizawa. Distribution of pores within alveoli in the human lung. Journal of Applied Physiology (Bethesda, Md.: 1985), 63(5):1866–1870, November 1987. ISSN 8750-7587. doi: 10.1152/jappl.1987.63.5.1866.

Mohammad F. Kiani and Antal G. Hudetz. A semi-empirical model of apparent blood viscosity as a function of vessel diameter and discharge hematocrit. Biorheology, 28(1-2):65–73, January 1991. ISSN 0006-355X. doi: 10.3233/BIR-1991-281-207. Publisher: IOS Press.

M. Matsuda, Y. C. Fung, and S. S. Sobin. Collagen and elastin fibers in human pulmonary alveolar mouths and ducts. Journal of Applied Physiology (Bethesda, Md.: 1985), 63(3):1185–1194, September 1987. ISSN 8750-7587. doi: 10.1152/jappl.1987.63.3.1185.

R. R. Mercer, M. L. Russell, and J. D. Crapo. Alveolar septal structure in different species. Journal of Applied Physiology (Bethesda, Md.: 1985), 77(3):1060–1066, September 1994. ISSN 8750-7587. doi: 10.1152/jappl.1994.77.3.1060.

Jordi Mill, Victor Agudelo, Chi Hion Li, Jérôme Noailly, Xavier Freixa, Oscar Camara, and Dabit Arzamendi. Patient-specific flow simulation analysis to predict device-related thrombosis in left atrial appendage occluders. REC Interventional Cardiology, 2021a. ISSN 2604-7322. doi: 10.24875/RECICE.M21000224. Accepted: 2022-07-04T12:28:00Z Publisher: S. Hirzel Verlag (SHV).

Jordi Mill, Josquin Harrison, Benoit Legghe, Andy L. Olivares, Xabier Morales, Jerome Noailly, Xavier Iriart, Hubert Cochet, Maxime Sermesant, and Oscar Camara. In-Silico Analysis of the Influence of Pulmonary Vein Configuration on Left Atrial Haemodynamics and Thrombus Formation in a Large Cohort. In Daniel B. Ennis, Luigi E. Perotti, and Vicky Y. Wang, editors, Functional Imaging and Modeling of the Heart, Lecture Notes in Computer Science, pages 605–616, Cham, 2021b. Springer International Publishing. ISBN 978- 3-030-78710-3.

Wolter Mooi and C. A. Wagenvoort. Decreased numbers of pulmonary blood vessels: Reality or artifact? The Journal of Pathology, 141(4):441–447, 1983. ISSN 1096-9896. doi: 10.1002/path.1711410403. _eprint: https://onlinelibrary.wiley.com/doi/pdf/10.1002/path.1711410403.

Christian Mühlfeld. Stereology and three-dimensional reconstructions to analyze the pulmonary vasculature. Histochemistry and Cell Biology, 156(2):83–93, August 2021. ISSN 1432-119X. doi: 10.1007/ s00418-021-02013-9. URL 10.1007/s00418-021-02013-9.

Christian Mühlfeld, Ewald R. Weibel, Ute Hahn, Wolfgang Kummer, Jens R. Nyengaard, and Matthias Ochs. Is Length an Appropriate Estimator to Characterize Pulmonary Alveolar Capillaries? A Critical Evaluation in the Human Lung. The Anatomical Record, 293(7):1270–1275, 2010. ISSN 1932-8494. doi: 10.1002/ar. 21158.

Y Nagasaka, J Bhattacharya, S Nanjo, M A Gropper, and N C Staub. Micropuncture measurement of lung microvascular pressure profile during hypoxia in cats. Circulation Research, 54(1):90–95, January 1984. doi: 10.1161/01.RES.54.1.90. Publisher: American Heart Association.

Matthias Ochs, Jens R. Nyengaard, Anja Jung, Lars Knudsen, Marion Voigt, Thorsten Wahlers, Joachim Richter, and Hans Jørgen G. Gundersen. The number of alveoli in the human lung. American Journal of Respiratory and Critical Care Medicine, 169(1):120–124, January 2004. ISSN 1073-449X. doi: 10.1164/ rccm.200308-1107OC.

Maria Isabel Pons, Jordi Mill, Alvaro Fernandez-Quilez, Andy L. Olivares, Etelvino Silva, Tom De Potter, and Oscar Camara. Joint Analysis of Morphological Parameters and In Silico Haemodynamics of the Left Atrial Appendage for Thrombogenic Risk Assessment. Journal of Interventional Cardiology, 2022:1–10, March 2022. ISSN 1540-8183, 0896-4327. doi: 10.1155/2022/9125224.

K. K. Pump. The Circulation in the Peripheral Parts of the Human Lung. Diseases of the Chest, 49(2): 119–129, February 1966. ISSN 0096-0217. doi: 10.1378/chest.49.2.119.

Dieter Schlosser, Ernst Heyse, and Heinz Bartels. Flow rate of erythrocytes in the capillaries of the lung. Journal of Applied Physiology, 20(1):110–112, January 1965. ISSN 8750-7587. doi: 10.1152/jappl.1965. 20.1.110. Publisher: American Physiological Society.

Kerstin Schmid, Andreas Knote, Alexander Mück, Keram Pfeiffer, Sebastian von Mammen, and Sabine C. Fischer. Interactive, Visual Simulation of a Spatio-Temporal Model of Gas Exchange in the Human Alveolus. Frontiers in Bioinformatics, 1, 2022. ISSN 2673-7647. URL https://www.frontiersin.org/article/ 10.3389/fbinf.2021.774300.

R. S. Sobin, H. M. Tremer, and Y. C. Fung. Morphometric basis of the sheet-flow concept of the pulmonary alveolar microcirculation in the cat. Circulation Research, 26(3):397–414, March 1970. ISSN 0009-7330. doi: 10.1161/01.res.26.3.397.

Hagit Stauber, Dan Waisman, Netanel Korin, and Josué Sznitman. Red blood cell dynamics in biomimetic microfluidic networks of pulmonary alveolar capillaries. Biomicrofluidics, 11(1):014103, January 2017. ISSN 1932-1058. doi: 10.1063/1.4973930.

K. C. Stone, R. R. Mercer, P. Gehr, B. Stockstill, and J. D. Crapo. Allometric relationships of cell numbers and size in the mammalian lung. American Journal of Respiratory Cell and Molecular Biology, 6(2):235–243, February 1992. ISSN 1044-1549. doi: 10.1165/ajrcmb/6.2.235.

Masahiro Toshima, Yuko Ohtani, and Osamu Ohtani. Three-dimensional architecture of elastin and collagen fiber networks in the human and rat lung. Archives of Histology and Cytology, 67(1):31–40, March 2004. ISSN 0914-9465. doi: 10.1679/aohc.67.31.

Alin-Florin Totorean, Sandor Ianos Bernad, Tiberiu Ciocan, Iuliana-Claudia Totorean, and Elena Silvia Bernad. Computational Fluid Dynamics Applications in Cardiovascular Medicine—from Medical Image-Based Modeling to Simulation: Numerical Analysis of Blood Flow in Abdominal Aorta. In Dia Zeidan, Lucy T. Zhang, Eric Goncalves Da Silva, and Jochen Merker, editors, Advances in Fluid Mechanics: Modelling and Simulations, Forum for Interdisciplinary Mathematics, pages 1–42. Springer Nature, Singapore, 2022. ISBN 978-981-19143-8-6.

Harun Tuğcu, Gülşin Canoğullari, Yıldırım Karslioğlu, Yasemin Günay Balci, Kubilay Uzuner, and Coşkun Yorulmaz. Comparison of the Effects of Two Different Fixation Methods on Lung Morphology: An Experimental Study. Turkiye Klinikleri Journal of Medical Sciences, 33(1):177–183, 2013. ISSN 1300-0292, 2146-9040. doi: 10.5336/medsci.2012-30068.

Ewald R. Weibel. Morphometric and stereological methods in respiratory physiology including fixation techniques. Techniques in respiratory physiology. Elsevier Scientific Publishers Ireland; Sole distributors for the U.S.A. and Canada, Elsevier Scientific Pub. Co., County Clare, Ireland, New York, N.Y., 1984. OCLC: 11013075.

Ewald R. Weibel, William J. Federspiel, Fabienne Fryder-Doffey, Connie C. W. Hsia, Martin König, Vilma Stalder-Navarro, and Ruth Vock. Morphometric model for pulmonary diffusing capacity I. Membrane diffusing capacity. Respiration Physiology, 93(2):125–149, August 1993. ISSN 0034-5687.

Pablo Zurita and Daniel E. Hurtado. Computational modeling of capillary perfusion and gas exchange in alveolar tissue. Computer Methods in Applied Mechanics and Engineering, 399:115418, September 2022. ISSN 0045-7825. doi: 10.1016/j.cma.2022.115418.

